# Prevalence, Production and the Role of *Staphylococcus aureus* Superantigens in Cystic Fibrosis Lung Disease

**DOI:** 10.1101/2025.05.07.652658

**Authors:** Ying Sun, Buqu Hu, Sahil Chhabra, Jade Tolentino, Nina Suzuki, Jesse Natarajan, Zachary Harris, Gail Stanley, Colleen Bianco, Paul Planet, Shardulendra P. Sherchand, Rajan Adhikari, Javad Aman, Xuchen Zhang, Jon Koff, Govindarajan Rajagopalan

## Abstract

Staphylococcus aureus (SA), the most common cystic fibrosis (CF) lung pathogen, is uniquely capable of producing superantigen (SAg) exotoxins, which are the most potent activators of the immune system. Although the proinflammatory roles of SA-SAgs is well-established, their role in the immunopathogenesis of CF lung disease is unexplored. Herein, we demonstrate that 60-80% of pediatric and adult CF SA isolates carried at least one SA-SAg gene, with the former harboring potent SA-SAgs (Staphylococcal enterotoxin A and B) more frequently (30-60%). Biofilms of clinical SA isolates readily produced biologically active SA-SAgs in artificial sputum medium and purified SA-SAgs retained their bioactivity in human CF sputum in vitro. Repeated intratracheal challenge with purified SA-SAgs induced a robust pulmonary inflammatory response in CF mouse models (βENAC and CFGC transgenic mice) expressing HLA-DR3 in a dose-dependent manner, with the low dose favoring a type 2 eosinophilic lung inflammatory response, and a high dose eliciting a type 1 inflammatory response with neutrophilic lung inflammation and higher mortality. In vivo neutralization of IFN-γ also promoted SA-SAg-driven type 2 inflammation. Intratracheal infection with sub-lethal dose of a clinical SA isolate producing SEB, but not the SEB-deficient mutant isogenic SA strain, also elicited an eosinophilic inflammatory response.

## INTRODUCTION

Cystic fibrosis (CF) is a common monogenic disorder, caused by mutations in the Cystic Fibrosis Transmembrane Conductance Regulator (CFTR) gene [1]. While CF affects multiple organ systems, chronic respiratory infections are the primary cause of morbidity among people with CF (pwCF) [2, 3]. Although CFTR modulator therapies have significantly improved the quality of life for pwCF [4, 5], they do not entirely eradicate microbial pathogens from the airways [6–9]. Thus, persistent bacterial infections remain as a major contributor to chronic airway inflammation and acute pulmonary exacerbations in pwCF.

Among the respiratory pathogens linked to CF lung disease, *Staphylococcus aureus* (SA) is not only the earliest colonizer of the airways but often the most prevalent bacterium responsible for chronic airway infections in pwCF. Its presence is often associated with poorer clinical outcomes [10–14]. Studies have shown that SA can persist in the lungs of CF patients even after modulator therapy, in contrast to other pathogens such as *Pseudomonas aeruginosa*, highlighting the problem [15, 16]. Notably, even in healthy individuals, colonization of the upper airways with SA begins at birth and continues throughout life, with approximately 20-50% of the normal adult population asymptomatically harboring SA either intermittently or chronically [17]. Given the inherent epithelial cellular dysfunctions and compromised immune defenses in pwCF, it is not surprising that SA is the most prevalent pathogen colonizing their airways, particularly among children aged 11-17 years, coinciding with a period of declining lung function [11, 18–20]. In this era of highly effective modulator therapies, SA continues to be a predominant pathogen well into adulthood [21].

In CF airways SA frequently forms biofilms which confers significant resistance to both antibiotic treatment and the host immune response, thereby facilitating persistent colonization [22–24]. Consequently, SA is not only the most prevalent pathogen in CF lungs but also one of the most challenging to eliminate. Furthermore, SA is distinctive among CF pathogens due to its capacity to produce a family of exotoxins known as superantigens (SAgs). Staphylococcal SAgs (SA-SAgs) are recognized as the most potent activators of the immune system and drivers of inflammation [25].

Unlike conventional antigens, SAgs bind directly to major histocompatibility complex (MHC) class II molecules outside of the peptide-binding groove without requiring processing by antigen-presenting cells (APCs). MHC class II-bound SAgs subsequently interact with only with a selected families of the variable regions of the T cell receptor (TCR) expressed on adaptive T cells, directly activating them without necessitating CD4 or CD8 co-receptor involvement or additional costimulation [25]. This results in robust polyclonal activation of 20-60% of adaptive CD4^+^ and CD8^+^ TCR^+^ T cells, as well as innate T cells, such as Natural Killer T (NKT) cells and mucosal-associated invariant T cells (MAITs) that express these TCR variable region families [26–28]. Recently, a SA-SAg has been shown to even activate γδ T cells [29], thus expanding the potential targets for SA-SAg.

While activated innate and adaptive T cells can directly induce tissue and cellular injury, they also promote inflammation through cytokine release. Thus, SA-SAgs are implicated in various acute and chronic inflammatory diseases [25]. Overall, SA is not only the most prevalent respiratory pathogen in pwCF that is often difficult to eradicate, a high frequency of both methicillin-sensitive and methicillin-resistant SA strains isolated from pwCF are capable of producing SA-SAg [30–32], which are the most potent activators of the immune system. Nonetheless, the role of SA-SAgs in the development of lung disease in CF remains largely unexplored and necessitates a systematic investigation.

To accurately model the pulmonary and systemic effects of SA-SAgs in CF, robust animal models that respond to SA-SAgs in a manner analogous to humans are essential. While mouse models of CF exist, SA-SAgs do not bind as effectively to murine MHC class II molecules as they do to human MHC (also known as Human Leukocyte Antigen or HLA) class II molecules [33–35]. Consequently, the pathological roles of SA-SAgs may not be fully appreciated in conventional mouse models [36, 37]. Ferret and porcine CF models have the same drawback [38]. We and others have demonstrated that transgenic expression of human MHC (HLA) class II molecules in mice renders them highly susceptible to the pathogenic effects of SA-SAgs and enables the study of many SA-SAg-induced diseases, including airway inflammation and lung injury in non-CF settings [28, 35, 39–43]. Therefore, to specifically investigate the role of SA-SAgs in pulmonary inflammation in the settings of CF, we developed two CF mouse models that also express HLA class II molecules. Our extensive studies using clinical CF SA isolates and the humanized mouse models provide novel insights into the previously underappreciated roles of SA-SAgs in the immunopathogenesis of lung disease in pwCF.

## MATERIALS AND METHODS

### Mice

CFTR ΔF508/ΔF508 “gut corrected” bitransgenic mice (generously provided by Dr. Craig Hodges, Case Western Reserve University) [44] and βENaC transgenic mice expressing the epithelial Na^+^ channel β subunit (βENaC protein, Scnn1b gene), driven by the Clara cell secretory protein promoter (a generous gift from Dr. Richard C. Boucher, University of North Carolina at Chapel Hill), both on a B6 background [45], were crossed with HLA-DR3 transgenic mice, also on the C57Bl/6 (B6) background [41], to generate HLA-DR3.CFGC and HLA-DR3.βENaC transgenic mice (hereafter referred to as HLA-DR3.ENAC mice), respectively. All mice were genotyped by PCR using genomic DNA as per standard technique. All mice were bred and maintained at the Yale University School of Medicine animal facility by the Division of Animal Care in Specific Pathogen-free facility following standard husbandry practices. All animal experiments were approved by Institutional Animal Care and use Committee.

### Generation of mutant *Staphylococcus aureus* (MNHOCH) lacking SEB

MNHOCH is a clinical methicillin-sensitive SA strain isolated from a patient with non-menstrual toxic shock syndrome (TSS) in the United States and produces high amounts of SEB. Mutant MNHOCH lacking SEB (MNHOCHΔSEB) was generated in IBT Bioservices (Rockville, MD). Electro-competent RN4220 cells were generated and stored at −80°C until use [46]. Approximately, 0.8 kb of sequence flanking SEB gene from the toxin harboring strain was cloned into pJB38 plasmid vector as described previously [47]. Electro-competent RN4220 cells were transformed using the plasmid, pJB38_SEB_Del. Donor cells transformed with plasmids were infected with phi11 phage. Phage lysates were collected and used for transduction of SA strain MNHOCH (pJB38_SEB_Del). Clones were selected by growing at 30°C in the presence of 10 ug/mL of Chloramphenicol. 1st allelic exchange and selection were done at 42°C and in the presence of 10 ug/mL of Chloramphenicol. 2nd allelic exchange selection is done at 30°C and clones sensitive to 10 ug/mL of Chloramphenicol were selected. Correct clones with gene knockouts were selected and confirmed by PCR using primers specific to the sequence flanking gene to be removed. Confirmation of SAg knockout is done by immunoblots (Supplementary Figure 1), sensitivity to 10 ug/mL of chloramphenicol, hemolytic typing, and sequencing of PCR products.

### Identification of SA-SAg gene distribution in CF SA isolates by multiplex PCR and by genomic DNA sequence data mining

Deidentified adult CF SA isolates were obtained from the Center for Phage Biology and Therapy at Yale, gown overnight in tryptic soy broth and bacterial DNA was extracted using DNA extraction kit (Qiagen). Multiplex PCR was performed using the genomic DNA as template to identify the presence of various SA-SAgs as described by Omoe et al., with minor modifications [48]. FemA and FemB genes were used as positive controls. In addition, we mined the whole genome DNA sequencing data of 104 pediatric CF SA isolates at Children’s hospital of Philadelphia for the distribution of various SA-SAg genes. Briefly, whole genome sequencing reads were obtained from NCBI Bioproject accession number PRJNA734908. Reads were trimmed and assembled using “shovill with the skesa assembler” option. Genomes were annotated using Prokka [49]. Toxin genes were found using “Abricate” utilizing a database containing toxin genes from various *S. aureus* strains (Supplemental file 1).

### CF sputum collection and processing

Spontaneously expectorated sputum samples collected from an adult pwCF negative for SA were mixed with PBS (three times the volume of sputum), vortexed vigorously, and homogenized by repeatedly passing through an 18-gauge needle (about 20 times). The mix was then clarified by centrifugation at 14000 rpm for 10 minutes and supernatants were stored frozen at −80°C. To check the stability of SA-SAgs in CF sputum, 10 μg of staphylococcal enterotoxin B (SEB, described below) was added to 1 ml of sputum extract collected as above or PBS and incubated at 37°C. The next day, mononuclear cells were prepared from spleens of HLA-DR3 transgenic mice and cultured with SEB (@ 1 μg/ml in RPMI) that had been incubated with CF sputum or PBS for 18 hours. The next day, the culture supernatants were collected and the amount of IL-2 in the supernatants was determined by ELISA (Biolegend, CA). For in vivo studies, HLA-DR3 transgenic mice were challenged with SEB that has been incubated with CF sputum extract or PBS as above (@ 10 μg/mouse) by intraperitoneal route. Three days later, spleens were collected and the distribution of TCR Vβ6^+^ and TCR Vβ8^+^ T cells within the CD4^+^ or CD8^+^ T cell subsets were determined by flow cytometry as described earlier [41].

### Production of SA-SAgs by CF SA biofilms grown in artificial sputum medium (ASM)

Single colonies of seven random CF isolates, MNHOCH (SEB^+^SA) and MNHOCHΔSEB (SEB^-^ SA), grown on tryptic soy agar plates, were inoculated into TSB and grown overnight at 37°C with shaking. The following day, 10 μl of the cultures was inoculated into 1 ml of ASM, (SCFM2. SynthBiome, GA) [50] in 24-well tissue culture plates. SCFM2 is a medium suitable for the growth of *Pseudomonas aeruginosa* and/or SA in an environment that mimics cystic fibrosis lung environment. This medium is supplemented with DNA, mucin, N-acetyl glucosamine, and dioleoylphosphatidylcholine in addition to other ions, free amino acids, glucose, and lactate to mirror CF sputum. The plates were incubated at 37°C without shaking to aid the formation of biofilm. The next day, 500 μl of the culture supernatants was harvested, spun at high speed for 10 minutes and the clear supernatants were stored at −80°C. The presence of SA-SAg in the biofilm culture supernatants was determined by HLA-DR3 splenocyte stimulation/cytokine production assay as described below.

Briefly, spleens from HLA-DR3 transgenic mice were collected aseptically and splenic mononuclear cells were prepared as per standard techniques. The cell concentration was adjusted to two million cells per ml in RPMI growth medium with serum and seeded in 24-well tissue culture plate with 1 ml in each well. One hundred microliters of the ASM bacterial biofilm culture supernatants prepared as above or 100 μl ASM alone were added to the splenocyte suspension. The negative control wells with splenocyte wells did not receive any ASM. As positive control, splenocytes were incubated with 1 μg of SEB. After overnight incubation, the splenocyte culture supernatants were collected and the concentration of IL-2 and IFN-γ in the culture supernatants were determined by ELISA (Biolegend, CA).

### Administration of staphylococcal enterotoxins

Lyophilized, endotoxin-reduced staphylococcal enterotoxins A and B (SEA and SEB) (Toxin Technologies, Sarasota, FL) were reconstituted in PBS and stored frozen in aliquots at −80°C. Equal amounts of highly purified SEC1, SEC2 and SEC3 were mixed to make SEC cocktail in PBS and stored frozen in aliquots at −80°C. Eight- to sixteen-week-old male and female mice were deeply sedated by isoflurane. Mice were challenged with 50 μl of SA-SAgs or PBS by oropharyngeal route on days 0, 3 and 6 and the animals were euthanized on day 8 to collect samples. In some experiments, anti-IFN-γ antibodies (Clone H22, BioXcell, 250 μg/mouse)) were given on day 0 by intraperitoneal route, immediately after intratracheal challenge with SEB.

### Intra-tracheal infection with live SA

A day prior to establishment of infection, single colonies of MNHOCH (SEB^+^SA) and its SEB-deletion mutant MNHOCHΔSEB (SEB^-^SA) were inoculated in TSB and grown overnight with shaking. The next day, small aliquots of the cultures were inoculated into fresh TSB at 1 in 10 dilution and grown till they reach a OD_600_ of 0.5 (usually 2-3 hours). Bacteria were washed twice with PBS by centrifugation and resuspended at a concentration of 10^6^ CFU in 50 μl PBS. The CFUs were confirmed on an aliquot of the inoculum by serial plating. Mice under isoflurane anesthesia were challenged with 50 μl of the bacterial suspension by oropharyngeal route and were closely monitored till the end of the experiment.

### Collection of Broncho-alveolar lavage (BAL) fluids

At indicated time points BAL fluids were collected by cannulating the exposed trachea and lavaging the lungs with 1 ml of PBS. BAL cells separated by centrifugation, and counted using an automated cell counter (Beckman) as described previously [51]. BAL protein concentration was determined using Pierce™ Bicinchoninic Acid (BCA) Protein Assay Kit (ThermoFisher). The concentrations of various cytokines and chemokines in the BAL were determined using multiplex kits (EMD Millipore).

### Preparation of Lung mononuclear cells

After transcardial perfusion with 10 ml of PBS, the right lobes of the lungs were collected, minced with a pair of scissors and mononuclear cells were extracted by collagenase and DNAse digestion as per standard procedure. Red blood cells were lysed by ammonium chloride lysis, cells were washed twice in medium containing fetal calf serum, counted using an automated cell counter and used for further flow cytometric analysis.

### Histopathology and immunohistochemistry (IHC)

After transcardial perfusion with 10 ml of PBS, the left lungs were removed, fixed overnight in buffered formalin followed by 90% ethanol, sectioned, and stained with H&E at the Yale Pathology Tissue Services. IHC with anti-CD3 and anti-CD19 antibodies was also performed at the same facility. Sections were evaluated using Nikon Elcipse microscope using a Nikon DS-Ri2 camera and representative images were acquired. Eosinophils in the tissue sections were identified by immunostaining with rat anti-mouse major basic protein (MBP) antibody (clone MT2-14.7.3, purchased from the laboratory of Elizabeth Jacobsen, Mayo Clinic, AZ) and Goat anti-Rat IgG (H+L) Cross-Adsorbed Secondary Antibody, Alexa Fluor™ 594 (Invitrogen) as per standard procedure.

### Flow cytometry

Flow cytometry was performed on BAL cells, lung MNCs and splenocytes to determine the distribution of various leukocyte subsets. Briefly, BAL cells, lung MNCs or splenocytes were first stained with fixable viability dye, followed by treatment with Fc block and then by fluorochrome conjugated antibodies diluted in Brilliant Stain Buffer (Thermo Fisher). The antibodies used are listed in Supp. Table 1. After staining with fluorochrome conjugated antibodies, the cells washed, were fixed in 0.1% formaldehyde for 30 mins, washed and resuspended in 300 μl of flow buffer. The next day, the cells were analyzed using a BD LSR II flow cytometer, when the simple myeloid panel was used or Aurora, spectral flow cytometer, when the larger lymphoid and myeloid panels were used. The flow cytometry gating strategy is outlined in Supp. Fig 2 [51, 52].

**Table 1.**
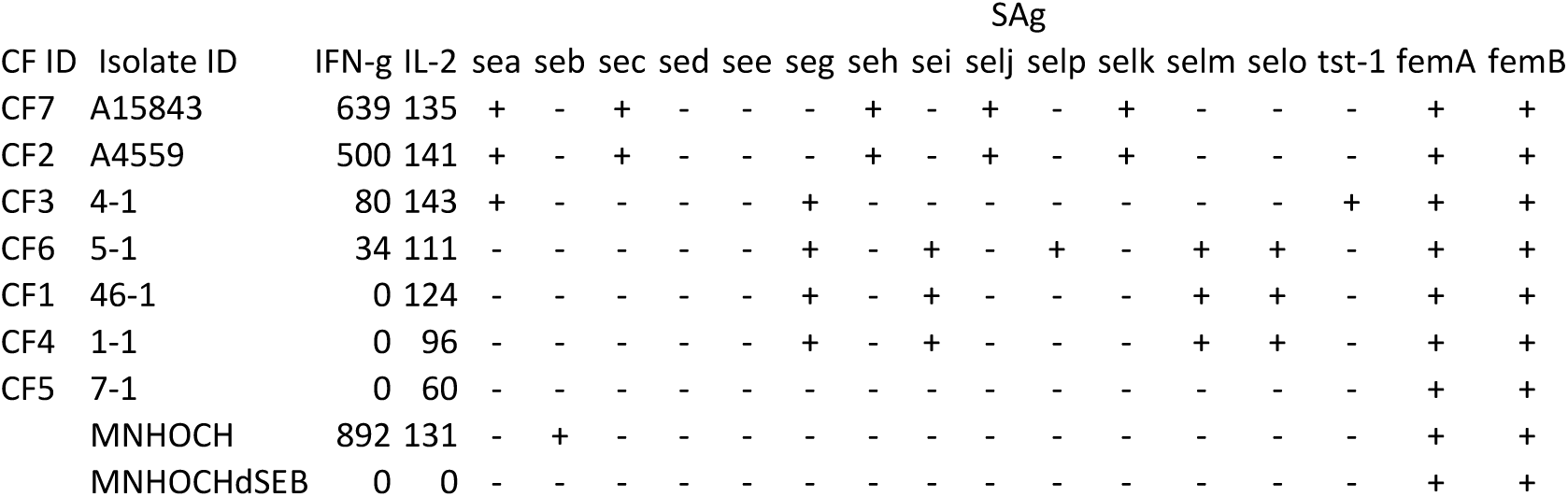
IL-2 and IFN-γ production by HLA-DR3 mice splenocytes following stimulation with culture supernatants from MNHOCH, MNHOCHΔSEB and clinical CF SA isolates (CF 1 – 7) grown as biofilms in artificial sputum medium.

### Statistics

Statistical significance of the data was determined using unpaired, non-parametric Mann-Whitney test or Student’s t test for two group comparisons. For comparison of survival curves, Log-rank (Mantel-Cox) test was used. All statistical calculations and generation of charts were performed using GraphPad Prism software (Prism 10).

## RESULTS

### Prevalence of SA-SAg genes in adult and pediatric CF SA isolates

In adult CF SA isolates, SEG was the most frequently identified SA-SAg (79%), followed by SEM (50%), SEO (46%), and SEN (38%) (Fig 1A). Among the SA-SAgs that are known to cause life-threatening systemic inflammatory conditions, SEA was more common (25%) than SEB or TSST-1 (8% each). We also assessed the number of SA-SAgs present in each isolate. Approximately 20% of the isolates contained at least three SA-SAgs, while 17% carried five or more (Fig 1B). In pediatric CF isolates, SEG was most prevalent (71%), followed by SEM, SEN, and SEO (63%) (Fig 1C). Interestingly, SEB was detected more frequently than SEA (59% vs. 32%) in this population, while the frequency of TSST-1 was only 7%. Notably, while 18% of the sequencing data did not identify any SA-SAg gene, over 50% of pediatric CF isolates carried five or more SA-SAg genes, with 19% containing up to eight different SA-SAg genes (Fig 1D). Overall, these findings underscore that (i) SA-SAg genes are frequently present in CF SA isolates, both pediatric and adult, and (ii) there are noteworthy differences in the distribution of SA-SAgs between pediatric and adult CF SA isolates.

**Figure 1.**
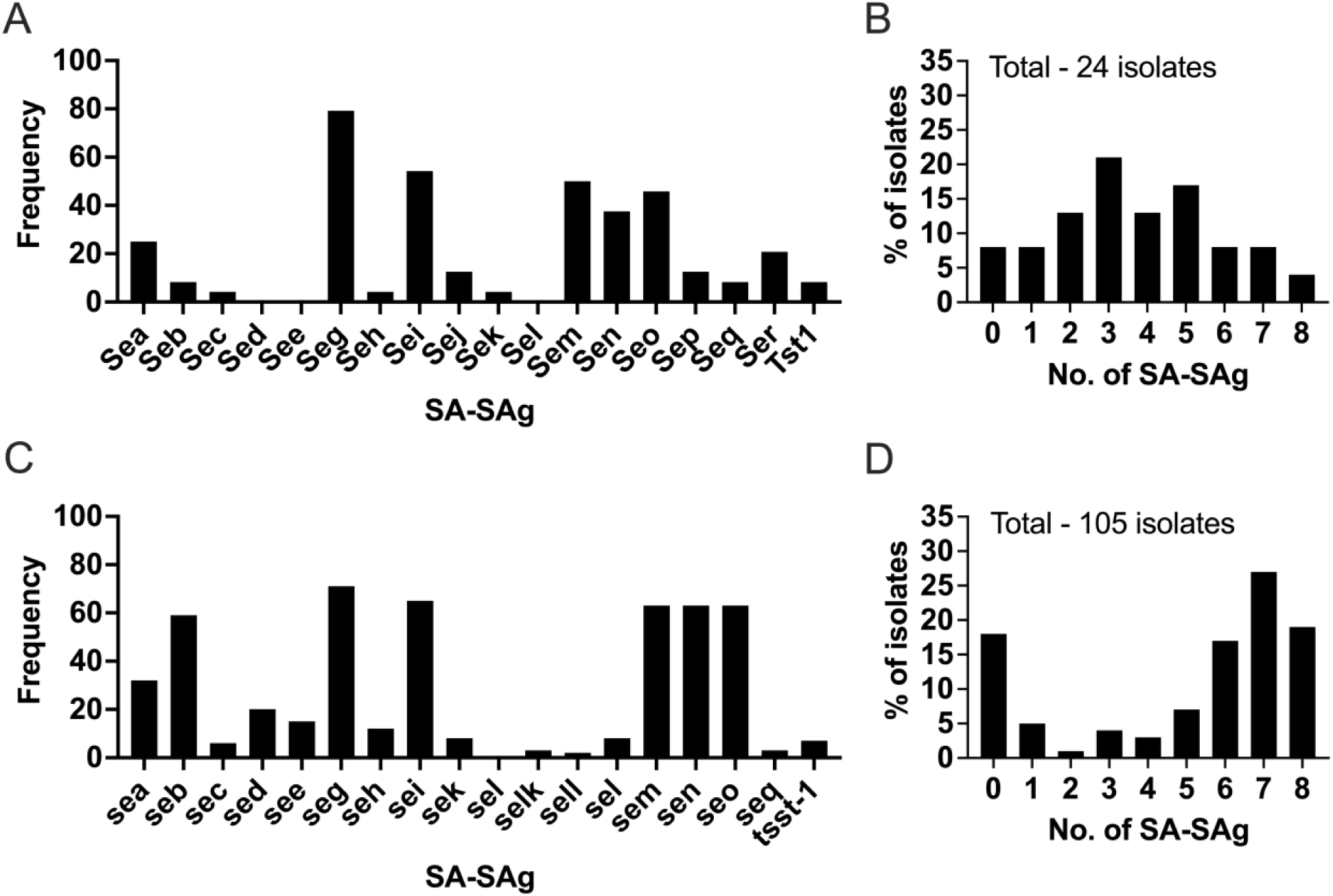
Distribution of SA-SAg genes in clinical CF isolates. (A and B) SA isolates from deidentified adult pwCF were grown in bacterial medium overnight and genomic DNA were column purified. The distribution of SA-SAg genes was determined by multiplex PCR. (A) The frequency of individual SA-SAg genes in all isolates and (B) distribution of SA-SAgs in each isolate. (C and D) Genomic sequencing data from pediatric CF SA isolates were analyzed for the distribution of SA-SAg genes. (C) The frequency of individual SAg genes in pediatric SA isolates and (D) distribution of SA-SAgs genes in individual isolates.

### Production of SA-SAgs by CF SA isolates in biofilm cultures in vitro

Next, seven random adult CF isolates were grown as biofilms in artificial sputum medium (ASM), and the presence of SA-SAgs in the biofilm supernatants was tested by in vitro cytokine production assay using HLA-DR3 splenocytes. Splenocyte culture supernatants incubated with ASM alone did not contain any IL-2 or IFN-γ indicating the absence of any potential mitogens in the ASM. Conversely, HLA-DR3 splenocyte culture supernatants incubated with purified staphylococcal enterotoxin B (SEB) contained very high concentrations of IL-2 and IFN-γ as expected (Table 1). Similarly, splenocytes cultured with supernatants from MNHOCH (SEB^+^SA) also contained very high levels of IL-2 and IFN-γ whereas addition of supernatants from MNHOCHΔSEB (SEB^-^SA) did not induce the production of any IL-2 or IFN-γ, confirming that supernatants from non-SA-SAg positive SA cultures does not contain any mitogens (Table 1). Interestingly, supernatants from all the seven CF clinical isolates induced the production of IL-2. However, supernatants from only 4 of these isolates induced the production of IFN-γ. Within these, there were significant differences in the concentration of IFN-γ produced, 2 isolates induced high levels of IFN-γ, one low and one intermediary levels of IFN-γ.

We next correlated the concentration of IL-2 nor IFN-γ with the SA-SAg profiles of these isolates. MNHOCHΔSEB (SEB^-^SA), which did not induce the production of IL-2 nor IFN-γ, did not harbor any of the SA-SAg tested (Table 1). MNHOCH (SEB^+^SA) was positive only for SEB. The two CF isolates (CF2 and 7), which induced high levels of IFN-γ (> 500 pg/ml) were positive for SEA and SEC, which are potent SA-SAgs. The isolate CF3, which induced intermediate levels of IFN-γ, was positive only for SEA. The isolate CF6, which induced very low levels of IFN-γ did not express SEA through SEE but was positive for 5 other SA-SAgs. Of the 3 isolates that failed to induce any IFN-γ, CF5 (that also induced the lowest levels of IL-2), did not express any of the SA-SAg tested, whereas the other 2 isolates were positive for 4 SA-SAgs. Overall, this data suggested that the expression of SEA through SEC correlated with strong induction of both IL-2 and IFN-γ, and the absence of these SA-SAgs and the presence of other SA-SAgs resulted in the production of only IL-2.

### SA-SAgs maintain their immunostimulatory activity in CF sputum

CF sputum exhibits distinct physicochemical properties compared to non-CF sputum, with higher viscosity, elasticity, and higher concentrations of mucins and proteases [53]. Most SA-SAgs are known enterotoxins that retain their biological activity in low gastric pH and resist various digestive proteases. Nonetheless, we next investigated whether the SA-SAgs retain their superantigenicity in chemically complex human CF sputum. SEB incubated with CF sputum extract elicited comparable IL-2 production from HLA-DR3 splenocytes as SEB incubated with PBS (Supp Fig 3A) and caused similar expansion of splenic CD4^+^ and CD8^+^ T cells expressing TCR Vβ8 in vivo in HLA-DR3 transgenic mice (Supp Fig 3B). These findings confirmed that SEB (and possibly all SA-SAgs), retain biological activity in human CF sputum.

### Intratracheal delivery of SEB induces a pulmonary inflammatory response

We challenged HLA-DR3.ENAC or B6.ENAC (mice expressing only endogenous mouse MHC class II molecules) via the intratracheal route, with 100 ng of SEB, a dose determined based on prior experience with non-CF pulmonary inflammation models [39] and studied a panel of parameters that reflect the extent of pulmonary inflammation.

First, BAL protein concentration was highest in HLA-DR3.ENAC mice challenged with SEB, significantly higher than that of HLA-DR3.ENAC mice challenged with PBS or B6.ENAC mice challenged with SEB (Fig 2A). However, BAL protein concentration did not significantly differ between B6.ENAC mice challenged with PBS or SEB. The total number of cells in the BAL exhibited a similar pattern (Fig 2B). Flow cytometric analysis of BAL cells revealed similar changes in various leukocyte subsets, except for eosinophils (Eos). AMs and different monocyte subsets were significantly higher in HLA-DR3.ENAC mice challenged with SEB compared to B6.ENAC mice challenged with SEB. PMNs followed a similar pattern.

**Figure 2.**
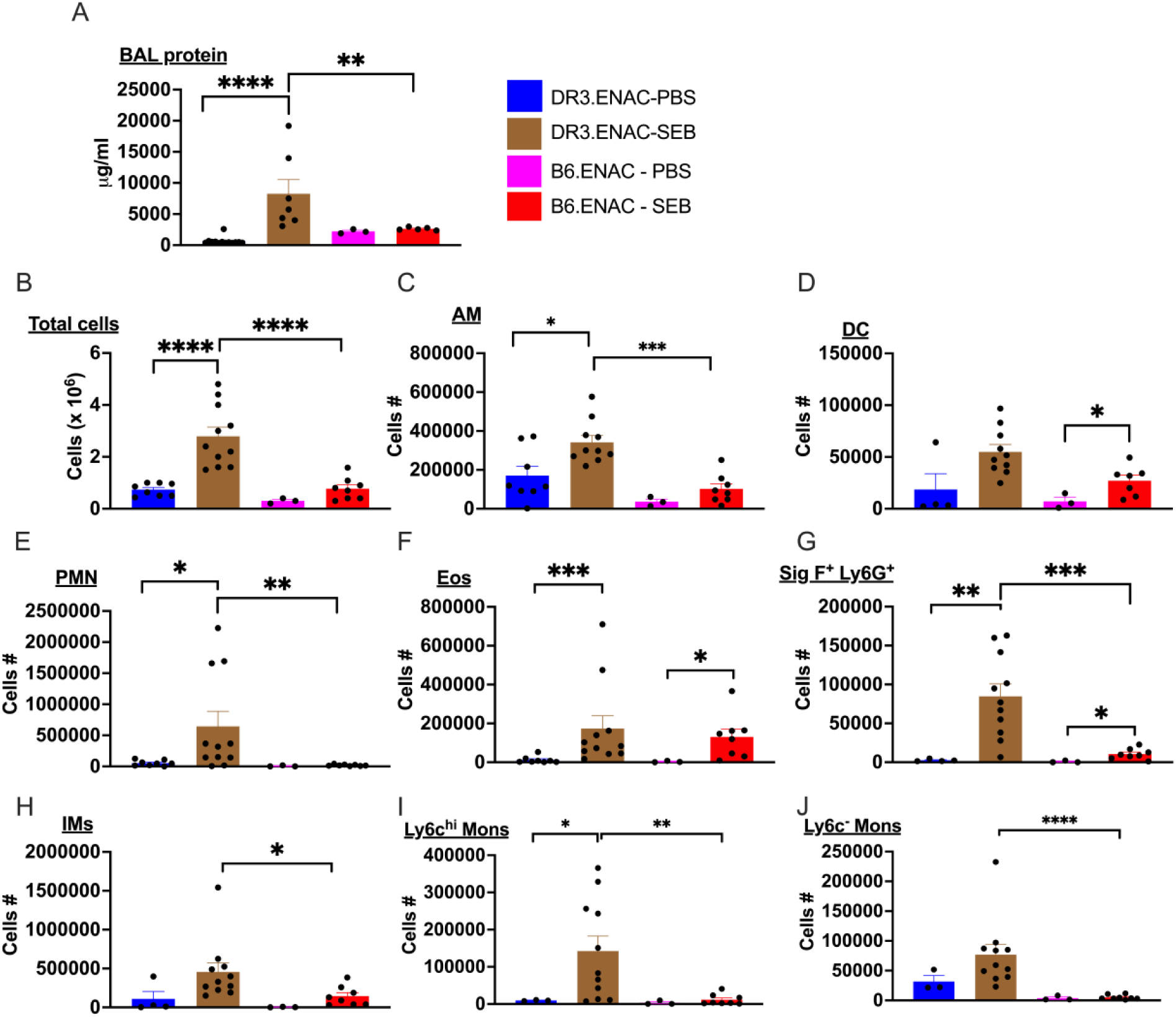
Repeated intratracheal administration of the staphylococcal superantigen, staphylococcal enterotoxin B (SEB), induces lung inflammation: HLA-DR3.ENAC and B6.ENAC mice were challenged intratracheally with PBS or 100 ng of SEB on days 0, 3 and 6 and killed on day 8. (A) BAL protein concentration as determined by BCA method. (B) Total cell counts as determined using a Coulter counter and (C through I) distribution of various BAL leukocyte subsets as determined by spectral flow cytometry. AM - alveolar macrophages, DC – Dendritic cells, PMN – Neutrophils, Eos – Eosinophils, IMs – interstitial macrophages. Data represents mean ± SE. * p<0.5, ** p<0.005, *** p<0.0005, **** p<0.00005.

Intratracheal administration of SEB consistently resulted in higher BAL Eos in both HLA-DR3.ENAC and B6.ENAC mice compared to PBS-challenged mice. While the mean Eos number was greater in HLA-DR3.ENAC mice treated with SEB than similarly treated B6.ENAC mice, it did not reach statistical significance (Fig 2F). The PMN:Eos was 3.7 for HLA-DR3.ENAC and 0.14 for B6.ENAC challenged with SEB, indicating a more robust neutrophilic response in HLA-DR3.ENAC mice. Notably, within the granulocyte gate, we also noted that the numbers of SigF^+^Ly6G^+^ cells were consistently elevated in SEB-challenged mice compared to PBS controls, with the highest levels observed in HLA-DR3.ENAC mice. Given the pronounced inflammatory response, we utilized HLA-DR3.ENAC mice in subsequent experiments, unless indicated.

Spectral flow cytometric analysis of CD45^+^ BAL cells from HLA-DR3.ENAC mice challenged with SEB revealed significant phenotypic differences in forward and side scatter profiles compared to PBS controls (Supp Fig 4). Additionally, there were notable changes in the expression patterns of CD11b and CD11c markers (Supp Fig 4). The scatter profiles of CD45^+^CD11b^+^CD11c^+^ cells (predominantly AMs) also differed in SEB-challenged mice. We further investigated the expression patterns of CD169, CD193, CD103, and CX3CR1 on various BAL cell types (Supp Fig 4). CD169 (Siglec1; sialoadhesin) is highly expressed on AM [54]. While AM from DR3.ENAC mice challenged with PBS expressed high levels of CD169, two distinct populations of CD169^+^ AM could be seen in mice challenged with SEB. A majority of the AMs from SEB-challenged mice downregulated CD169, while a small subset retained the high expression of CD169.

CD103, which is considered a marker for lung DCs, was expressed at higher levels on AMs and DCs from mice challenged with SEB. While the expression of CX3CR1 was not different on AMs, DCs from SEB-challenged mice expressed lower levels of CX3CR1. CD193 (CCR3) is the receptor for the chemokine eotaxin, which is responsible for recruitment of eosinophils to the lungs [55]. Compared to PBS-treated mice, the expression of CD193 on Eos and SigF^+^Ly6G^+^ cells from SEB-challenged mice was much lower. Overall, these results indicated that intratracheal SEB administration led to a significant increase in lung recruitment of various inflammatory cell types with distinct phenotypic changes compared to mice challenged with PBS.

Next, we assessed the distribution of various leukocyte subsets in the digested lungs of HLA-DR3.ENAC mice (Fig 3). Among the myeloid cell subsets, all cells, except conventional monocytes (cMons) and Eos, were significantly elevated in the lungs of SEB-challenged mice compared to those challenged with PBS. While cMons and Eos were higher in SEB-challenged mice, this did not reach statistical significance. Within the lymphoid population, total CD4^+^ and CD8^+^ T cells, CD4^+^ and CD8^+^ T cells expressing TCR Vβ6 and Vβ8, NKg2D^+^ cells and NK1.1^+^ cells were all significantly higher in SEB-challenged mice. The total numbers of γδ T cells were also significantly higher in SEB-challenged mice. It should be noted that γδ T cells are not considered to be reactive to SEB. Greater lung recruitment of γδ T cells indicated that the SEB-induced inflammatory response resulted in recruitment of many cell types into the lungs. Overall, these findings suggested that repeated intratracheal exposure to SEB elicited a robust pulmonary inflammatory response in both B6.ENAC and HLA-DR3.ENAC mice, with more pronounced changes in mice expressing HLA-DR3, corroborating previous observations.

**Figure 3.**
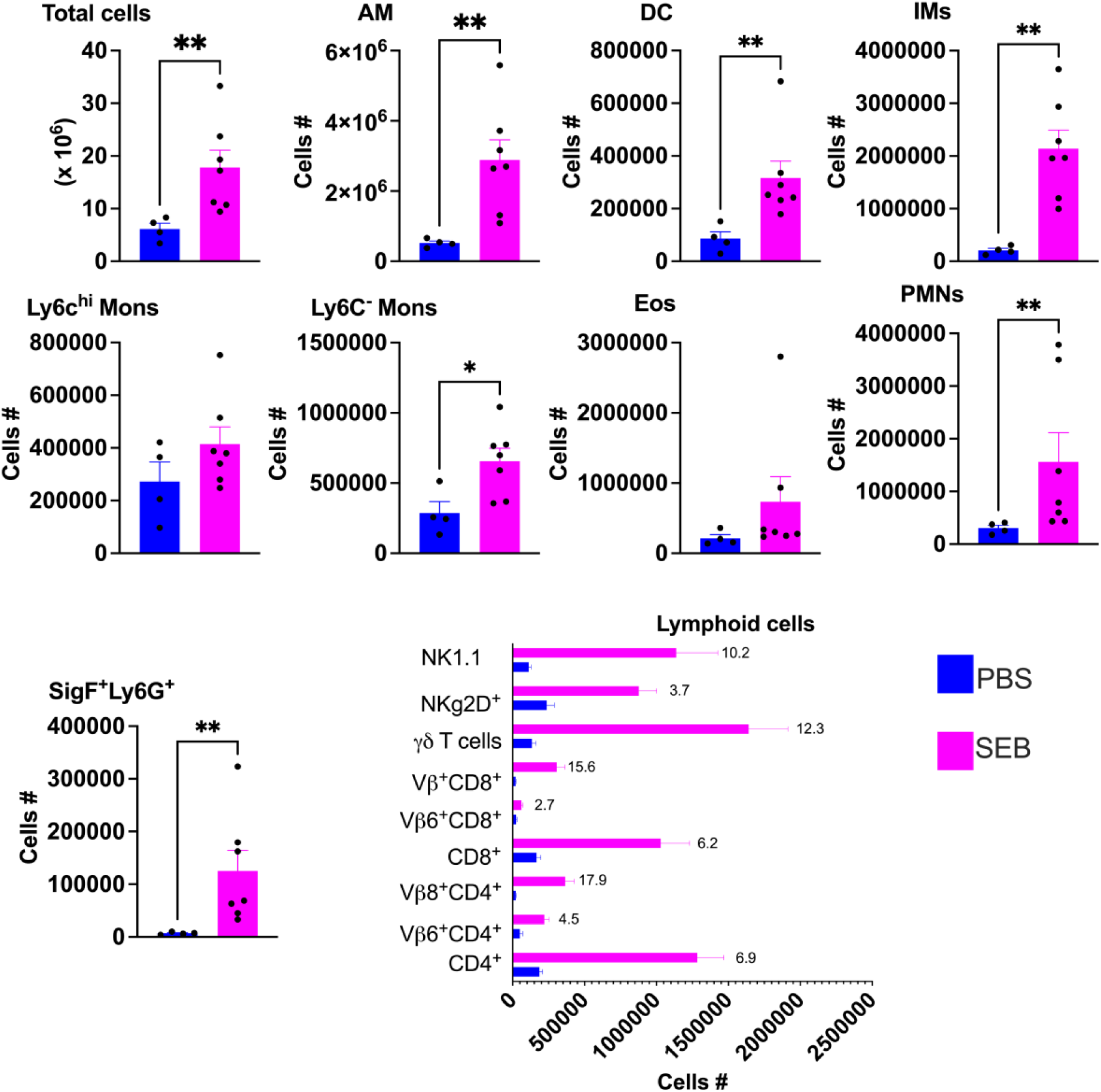
Lung recruitment of various leukocyte subsets following repeated intratracheal administration of SEB: HLA-DR3.ENAC mice were challenged intratracheally with PBS or 100 ng of SEB on days 0, 3 and 6 and killed on day 8. Distribution of various leukocyte subsets isolated from perfused digested lungs was determined by flow cytometry. AM - alveolar macrophages, DC – Dendritic cells, PMN – Neutrophils, Eos – Eosinophils, IMs – interstitial macrophages. Data represents mean ± SE. * p<0.5, ** p<0.005, *** p<0.0005, **** p<0.00005.

### Histopathology

As all lungs were lavaged (for cytokines and cell analysis) prior to harvesting lungs, obstruction of the major airways with mucus plugs was not appreciable. Lungs from mice challenged with PBS did not reveal major inflammatory changes except for mild peribronchial infiltration (Fig 4). However, lungs from HLA-DR3.ENAC mice challenged with SEB showed more severe inflammatory changes with extensive perivascular and peribronchial inflammation compared to B6.ENAC mice challenged with SEB. Eosinophilic deposits could be observed in the lungs of HLA-DR3.ENAC mice challenged with SEB, indicating mucous accumulation in distal/terminal airways/alveoli (Fig 4H). Moreover, some lung sections from HLA-DR3.ENAC mice revealed histopathological changes resembling organizing pneumonia (Fig 4 I and J)

**Figure 4.**
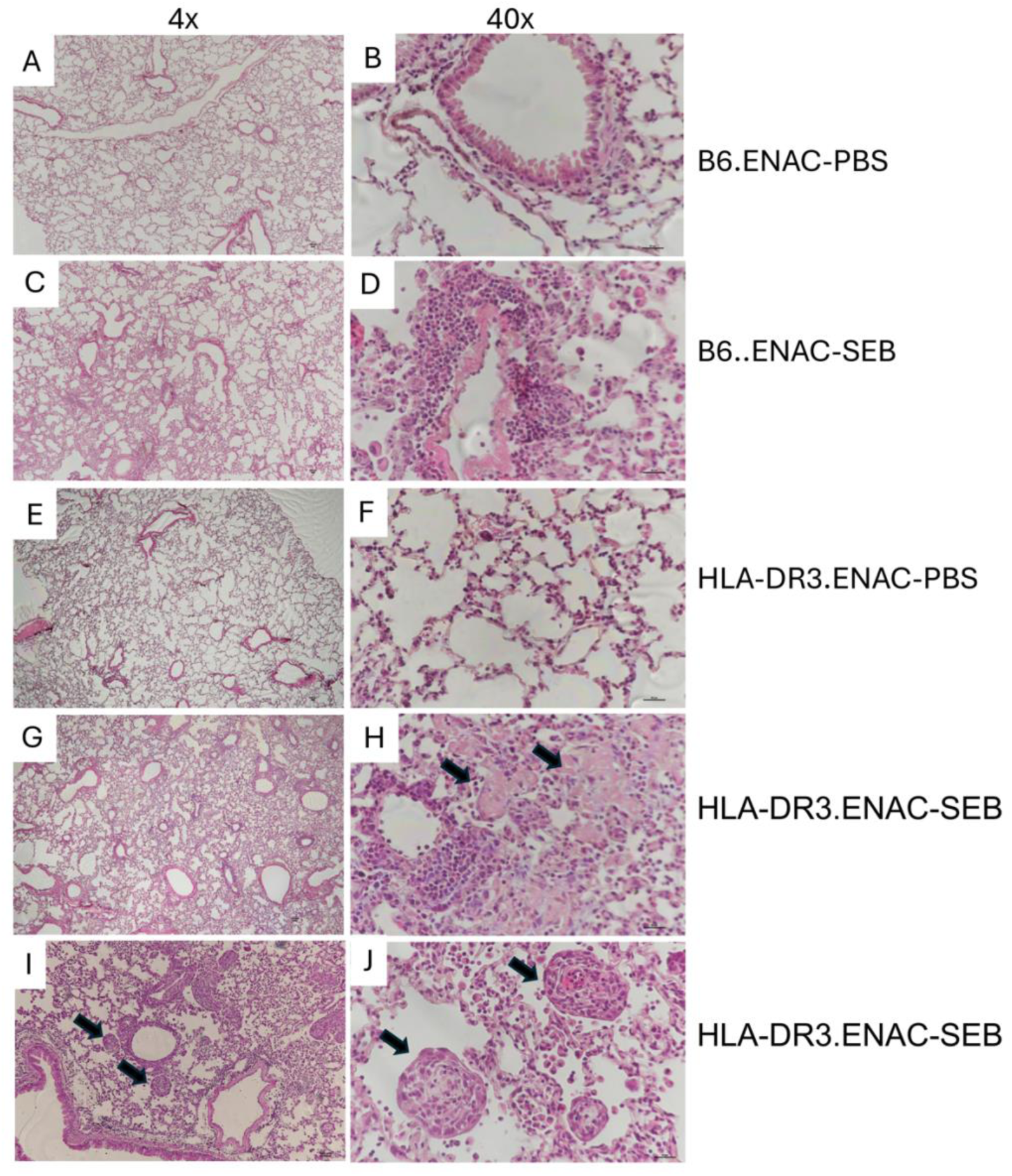
Repeated intratracheal administration of staphylococcal enterotoxin B (SEB) induces distinct lung inflammation pattern in HLA-DR3 .ENAC and B6.ENAC mice. HLA-DR3.ENAC mice were challenged intratracheally with PBS or 100 ng of SEB on days 0, 3 and 6 and killed on day 8. After collecting BAL, lungs were perfused, the right lobes were formalin fixed, sectioned and stained with hematoxylin and eosin. Representative photomicrographs at different magnifications. Arrows in panel H point to eosinophilic deposits, and in panels I and J point to organizing pneumonia-like features.

As lungs from mice challenged with SEB had significantly higher innate as well as adaptive T cells as determined by flow cytometry, we then analyzed the distribution of T cells among the inflammatory infiltrates by CD3 immunohistochemical staining. As can be seen from Supp. Fig 5, CD3^+^ cells were sparsely found in the lungs from PBS challenged ENAC mice. While SEB-challenged B6.ENAC mice did contain some CD3^+^ cells within the inflammatory infiltrates, they were highest in HLA-DR3.ENAC mice. The CD3^+^ cells were predominantly present in both peribronchial as well as peribronchial infiltrates. CD19^+^ cells could also be detected in the lungs of HLA-DR3.ENAC mice challenged with SEB, but not PBS. Clusters of CD19^+^ cells were present within the peribronchial as well as perivascular infiltrates (Supp. Figure 5). Immunostaining with anti-MBP antibodies also showed the widespread presence of Eos in the inflammatory infiltrates (Supp. Figure 5).

Next, we evaluated the extent to which transgenic expression of ENAC contributed to pulmonary inflammation. For this, wildtype HLA-DR3 transgenic mice without ENAC (HLA-DR3.WT) were challenged with SEB (100 ng) or PBS. Intratracheal exposure to SEB resulted in a significant increase in BAL PMNs and Eos even in HLA-DR3.WT mice (Fig 5). However, this was less pronounced compared to HLA-DR3 mice expressing ENAC. This is expected as HLA-DR3.ENAC mice challenged with PBS alone had a higher BAL total cell counts compared to HLA-DR3 mice lacking ENAC challenged with PBS (Fig 5A). It has been demonstrated previously that transgenic expression of ENAC per se can result in inflammation in the absence of exogenous insults [45]. SEA, another potent SA-SAg, also elicited inflammatory changes, albeit less severe than SEB (Supp. Fig 6).

**Figure 5.**
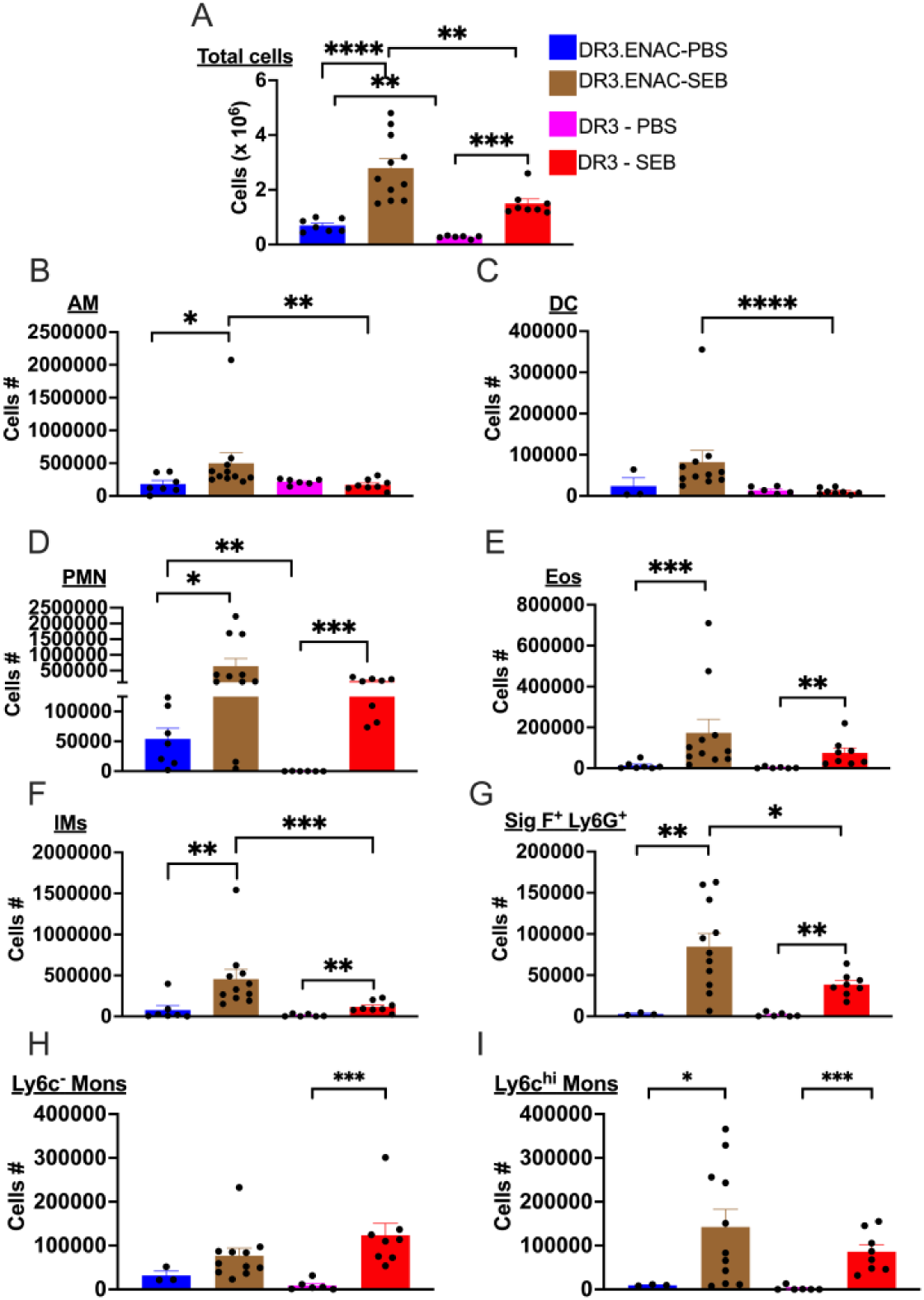
The impact of CF-like environment on the nature of lung inflammatory response following repeated intratracheal administration of SEB: HLA-DR3 transgenic mice expressing or not expressing ENAC were challenged intratracheally with PBS or 100 ng of SEB on days 0, 3 and 6 and killed on day 8. Distribution of various BAL leukocyte subsets was determined by flow cytometry. AM - alveolar macrophages, DC – Dendritic cells, PMN – Neutrophils, Eos – Eosinophils, IMs – interstitial macrophages. Data represents mean ± SE. * p<0.5, ** p<0.005, *** p<0.0005, **** p<0.00005.

We then confirmed the abilities of purified SA-SAgs to induce lung disease in another mouse model of CF namely, CFTR ΔF508/ΔF508 “gut corrected” mice (CFGC mice) expressing HLA-DR3. As shown in Supp. Fig 7, intratracheal administration of SEA, SEB and SEC all induced significant inflammatory cell recruitment to the lungs. However, SEA was least inflammatory whereas SEB and SEC were comparable. However, the overall inflammatory changes in CFGC mice were less severe than ENAC mice, which has been described earlier. These findings confirm our hypothesis that airway exposure to SA-SAgs can induce pulmonary inflammation. The extent of inflammatory response may be determined by the SA-SAg.

### Dose-dependent nature of pulmonary inflammatory responses elicited by SA-SAg

We next investigated whether the dose of SEB influenced the nature of the inflammatory response. HLA-DR3.ENAC mice were challenged with either 10 or 1000 ng of SEB as outlined in the methods. We observed significantly higher mortality in mice receiving 1000 ng of SEB compared to those receiving 10 ng (or even 100 ng) of SEB (Fig 6A). Based on these observations, we then challenged a group of B6.ENAC mice with 1000 ng of SEB and none of the B6.ENAC mice died in this cohort (0/15). This suggested that 1000 ng of SEB likely elicited a more robust pulmonary and systemic inflammatory response in HLA-DR3.ENAC mice resulting in higher mortality compared to B6.ENAC mice, reinforcing earlier findings regarding systemic toxicity with high-dose SEB [41, 56].

**Figure 6.**
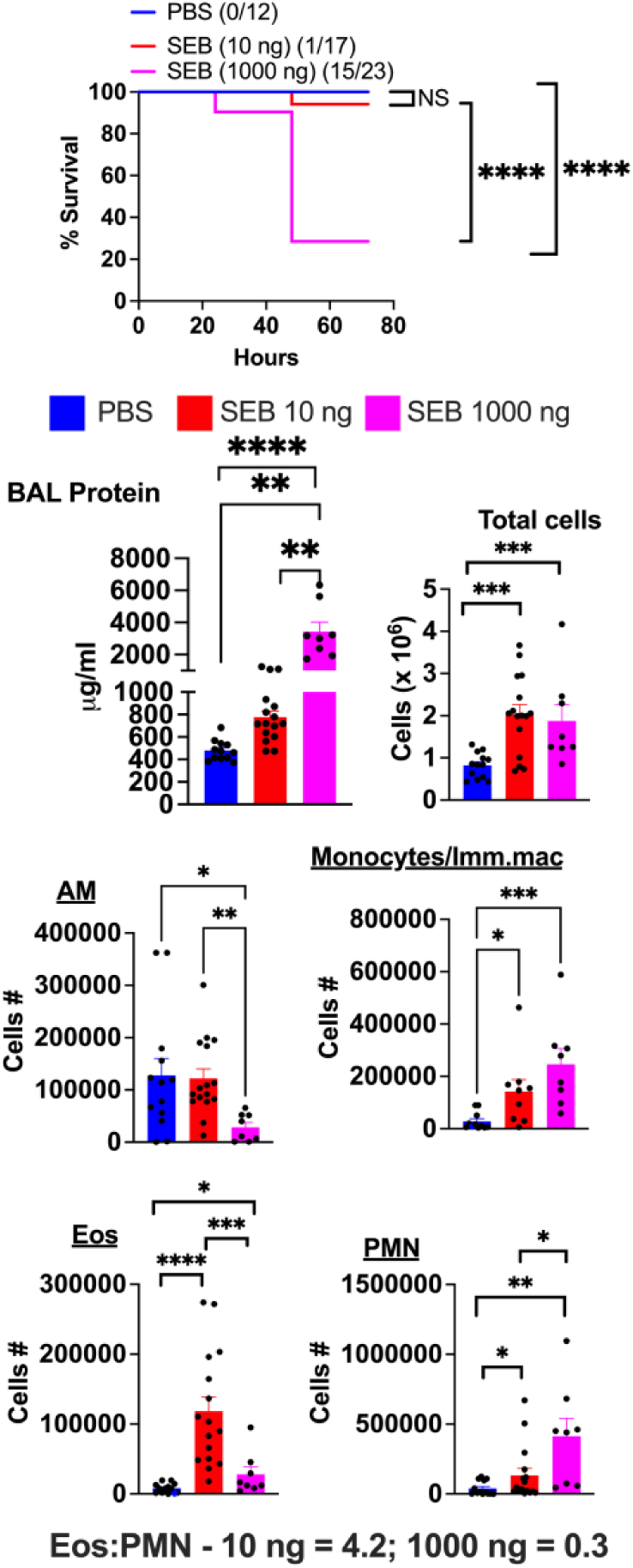
Repeated intratracheal administration of SEB induces a differential lung immune cell recruitment in a dose-dependent manner: HLA-DR3.ENAC mice were challenged intratracheally with PBS, 10 ng or 1000 ng of SEB on days 0, 2 and 4 and killed on day 6. (A) Mortality (B) BAL protein concentration (C) Total BAL fluid cell counts, and (D to F) distribution of various BAL leukocyte subsets by flow cytometry. Data represents mean ± SE. * p<0.5, ** p<0.005, *** p<0.0005, **** p<0.00005.

Regardless of the concentration, SEB consistently elicited a robust pulmonary inflammatory response, as evidenced by elevated BAL protein concentration (Fig 6B) and higher total BAL cell counts (Fig 6C). While total BAL cells were increased in all HLA-DR3.ENAC mice challenged with SEB, there was no significant difference between the 10 and 1000 ng groups. However, BAL protein concentration was highest in mice challenged with 1000 ng of SEB, followed by those receiving 10 ng. Importantly, the dose of SEB influenced the numbers of PMNs and Eos in the BAL. Mice challenged with 10 ng of SEB exhibited similar PMN and Eos counts in the BAL (133720 ± 48931 and 118428 ± 20488, mean ± SE, for PMNs and Eos, respectively; PMN:Eos = 1.13). In contrast, in mice challenged with 1000 ng of SEB, PMN numbers increased (413478 ± 128091), while Eos numbers decreased (27921 ± 10703), resulting in a PMN:Eos ratio of 14.1.

Histologically, lungs from mice challenged with 10 ng of SEB showed inflammatory changes with perivascular and peribronchial inflammation (Suppl. Fig 8). However, lung sections from mice challenged with 1000 ng of SEB showed no such distinctive perivascular or peribronchial infiltration, rather showed diffuse inflammation with hemorrhage and fibrinous deposits easily appreciated under higher magnification (Suppl. Fig 8). Next, we analyzed the concentrations of various cytokines and chemokines in the BAL of HLA-DR3.ENAC mice collected at the time of sacrifice (day 8) (Fig 7). The concentrations of several cytokines and chemokines were significantly elevated in the BAL of mice receiving 1000 ng of SEB compared to those challenged with PBS or 10 ng of SEB (Fig 7). Notably, IL-5, IL-13, IL-17, and IFN-γ were significantly higher in this group, correlating with the greater recruitment of both Eos and PMNs. Overall, a higher dose of SEB induced a more robust neutrophilic inflammation with increased mortality in HLA-DR3.ENAC mice, while a lower dose failed to recruit more PMNs to the lungs, leading to an eosinophil-predominant response in HLA-DR3.ENAC mice with minimal mortality.

**Figure 7.**
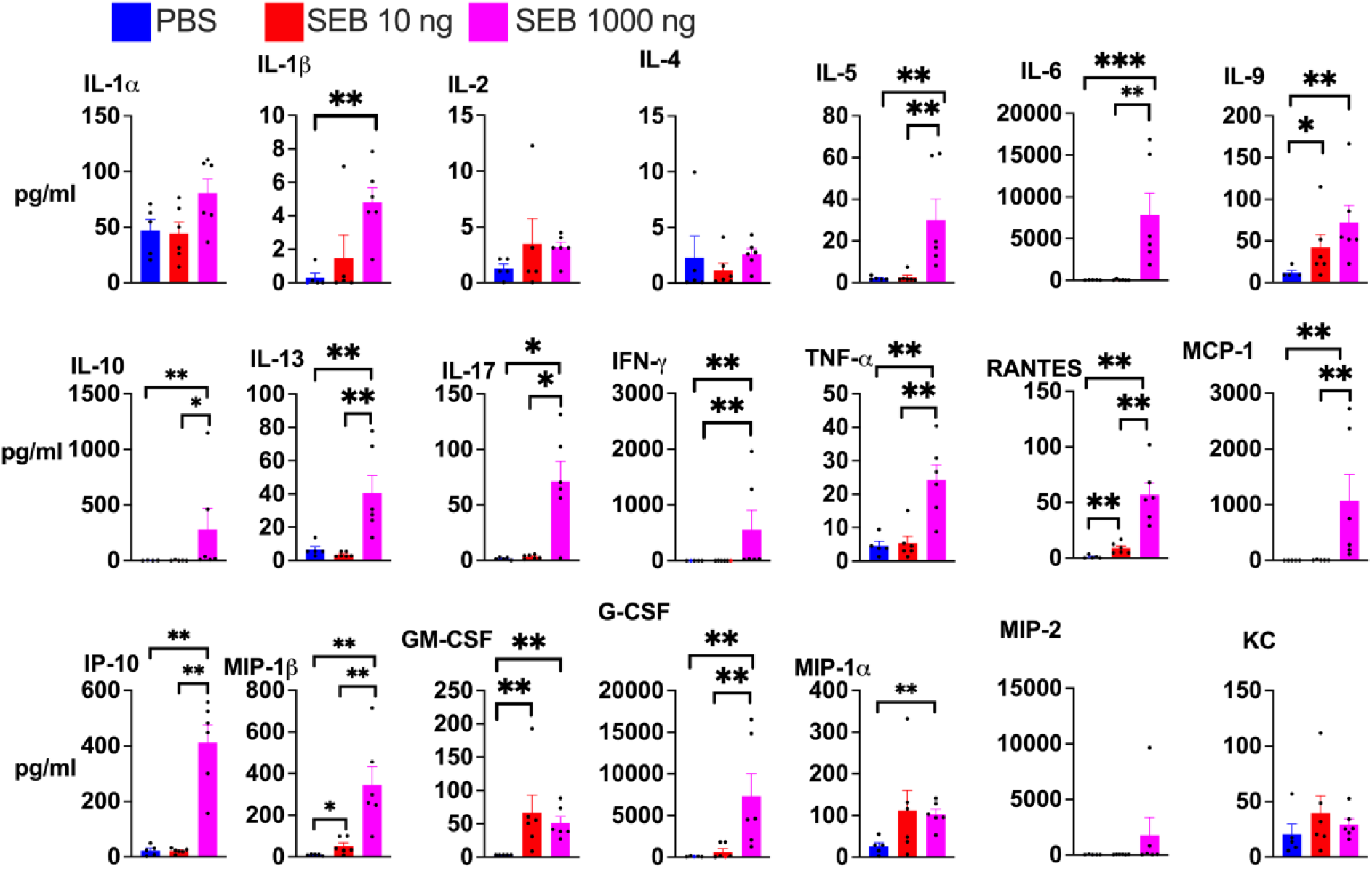
Cytokine concentrations in the BAL: HLA-DR3.ENAC mice were challenged intratracheally with PBS, 10 ng or 1000 ng of SEB on days 0, 2 and 4 and killed on day 6. The concentrations of various cytokines and chemokines in the BAL fluid was determined by multiplex assay. Data represents mean ± SE. * p<0.5, ** p<0.005, *** p<0.0005, **** p<0.00005.

### Effect of in vivo neutralization of IFN-γ on SEB-induced pulmonary inflammation

We recently demonstrated that neutralization of IFN-γ significantly increased lung recruitment of Eos and PMNs in a model of acute SEB-induced extrapulmonary acute lung injury in HLA-DR3 transgenic mice [51]. Similarly, it has been extensively that in vivo neutralization of IFN-γ promoted a type 2 eosinophilic response in several models of pulmonary inflammation [57–59]. Therefore, we investigated the effect of IFN-γ neutralization on SEB-induced pulmonary inflammation in CF settings, which has not been investigated before. In vivo neutralization of IFN-γ resulted in significantly higher Eos counts in the BAL of HLA-DR3.ENAC mice (Fig 8). The numbers of Sig F^+^Ly6G^+^ cells were also significantly elevated in these mice compared to HLA-DR3.ENAC mice challenged with SEB alone. Surprisingly, in SEB challenged B6.ENAC mice treated with anti-IFN-γ antibody, total BAL cell counts, including AM, DC, Eos, IMs, cMons, and Ly6C Mons, were all significantly elevated compared to SEB challenged mice not treated with anti-IFN-γ antibody. Notably, total Eos counts in B6.ENAC mice treated with anti-IFN-γ antibody were significantly higher than those in similarly treated HLA-DR3.ENAC mice.

**Figure 8.**
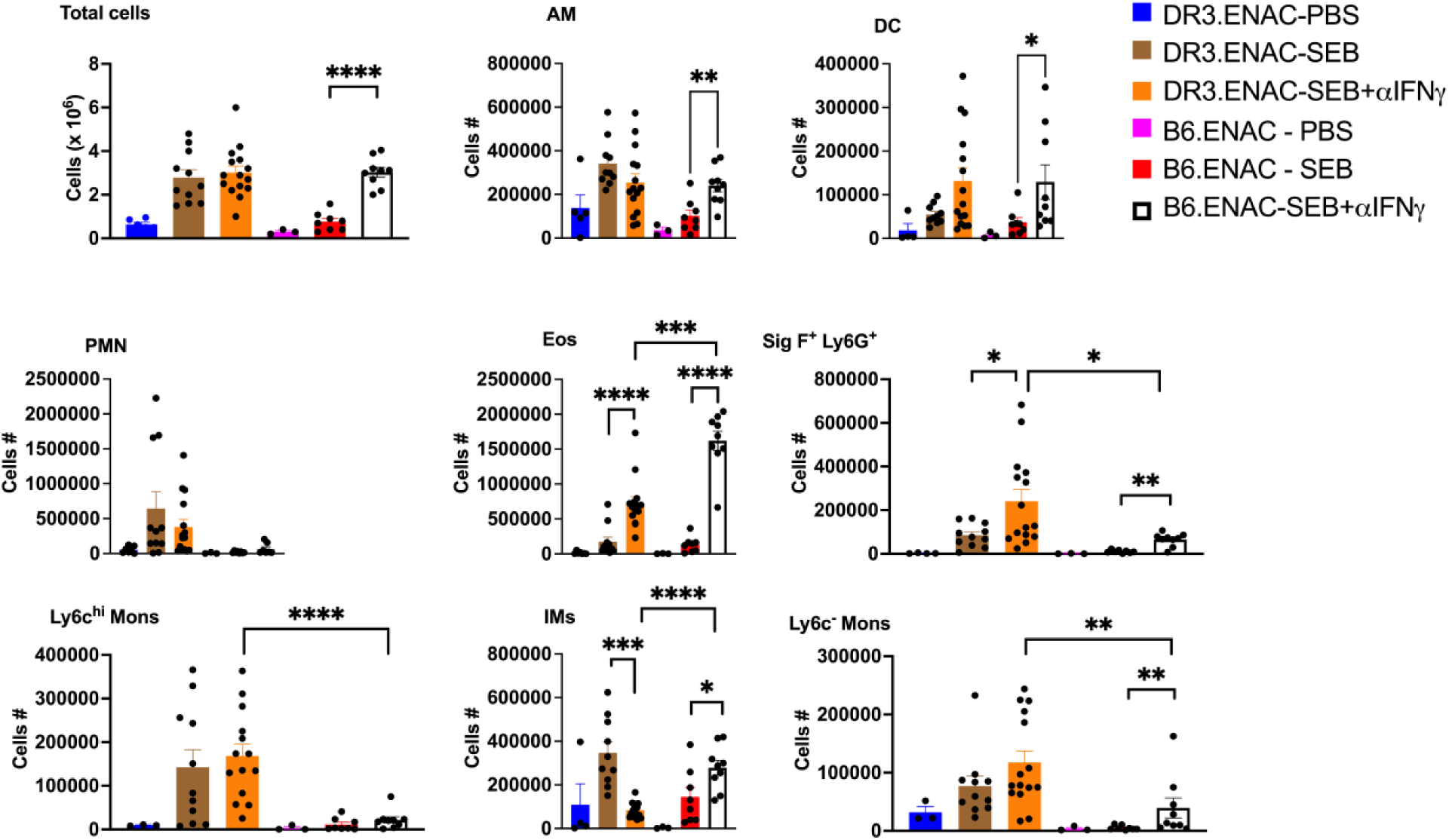
In vivo neutralization of IFN-γ alters the nature of immune cell infiltration in SEB-induced CF lung disease: HLA-DR3.ENAC or B6.ENAC mice were challenged intratracheally with PBS, 100 ng of SEB on days 0, 3 and 6 and killed on day 8. Neutralizing anti-IFN-γ antibodies (250 μg/mouse, IP route) were administered on day 0. Distribution of various BAL leukocyte subsets was analyzed by flow cytometry. AM - alveolar macrophages, DC – Dendritic cells, PMN – Neutrophils, Eos – Eosinophils, IMs – interstitial macrophages. Data represents mean ± SE. * p<0.5, ** p<0.005, *** p<0.0005, **** p<0.00005.

We next studied the concentrations of BAL cytokines and chemokines from DR3.ENAC and B6.ENAC challenged with SEB and treated or not treated with neutralizing anti-IFN-γ antibody. First of all, as shown in Fig 9, the concentrations of various cytokines such as G-CSF, GM-CSF, IL-1β, IL-6, IL-12p70, IL-17A, TNF-α, MIP-1α, MIP-1β, MCP-1 and more importantly IFN-γ and IP-10, a cytokine strongly induced by IFN-γ, were significantly higher in the BAL of DR3.ENAC mice challenged with 100 ng of SEB compared to B6.ENAC mice challenged with SEB underscoring a more robust type 1/3 response in HLA-DR3 transgenic mice. The BAL concentrations of almost all the cytokines/chemokines were less than that of DR3.ENAC mice challenged with 1000 ng of SEB and more than that of mice challenged with 10 ng of SEB, indicating a dose dependent effect. Treatment with neutralizing anti-IFN-γ antibody resulted in a significant reduction in the concentrations of most these cytokines in the BAL in HLA-DR3 ENAC mice. As the concentrations of most cytokines were already low or undetectably in BAL from B6.ENAC mice, the effect of anti-IFN-γ antibody could not be appreciated. However, the concentration of IL-4 was significantly higher in B6.ENAC mice treated with anti-IFN-γ antibody than similarly treated HLA-DR3.ENAC mice. Similarly, the concentration of IL-5 remained high in HLA-DR3.ENAC and B6.ENAC mice treated with anti-IFN-γ antibody (Fig 9). Overall, elevated type 2 cytokines (IL-4 and IL-5), correlated with elevated BAL Eos counts in anti-IFN-γ antibody-treated B6.ENAC mice.

**Figure 9.**
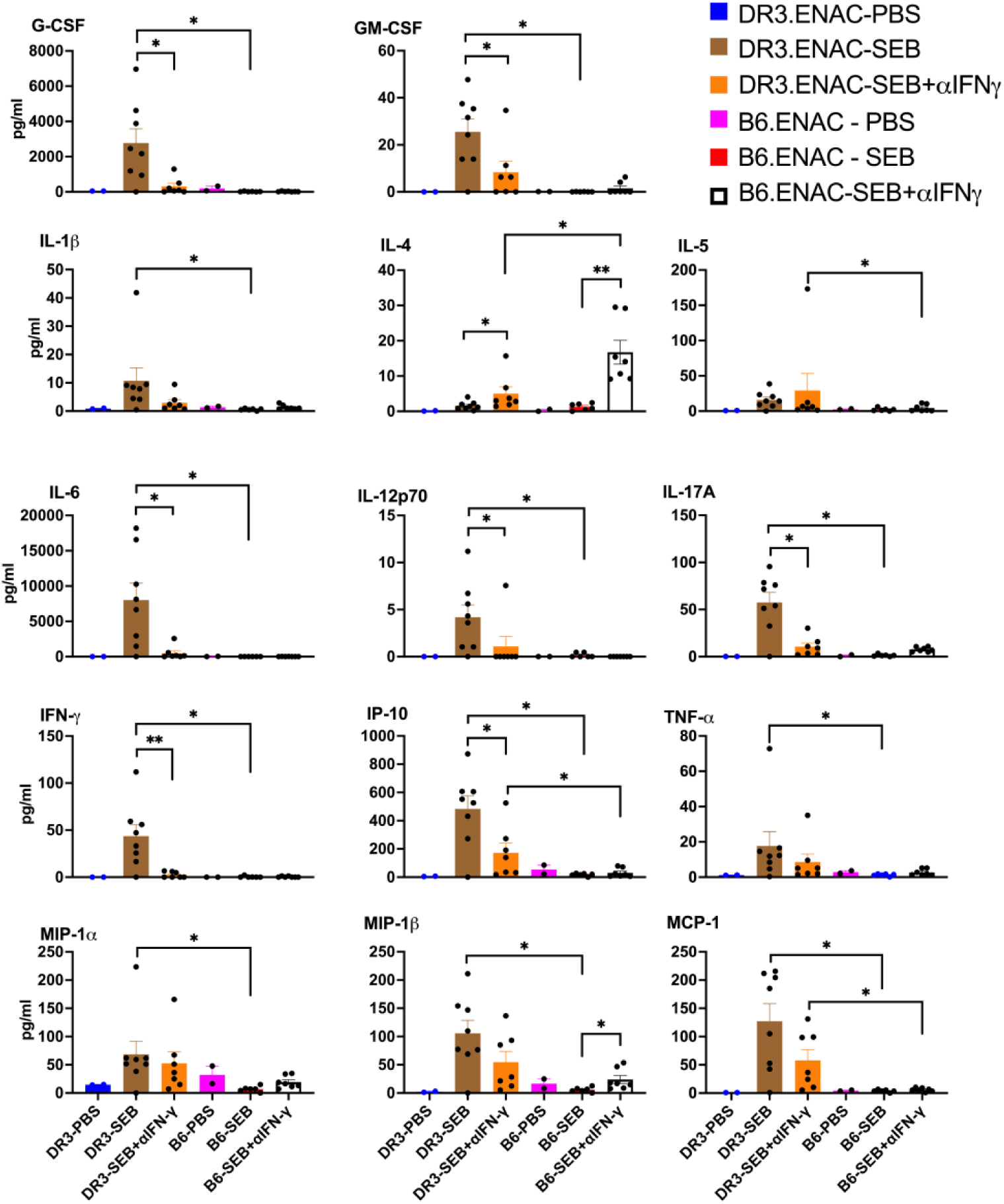
In vivo neutralization of IFN-γ alters the BAL cytokine profile: HLA-DR3.ENAC or B6.ENAC mice were challenged intratracheally with PBS, 100 ng of SEB on days 0, 3 and 6 and killed on day 8. Neutralizing anti-IFN-γ antibodies (250 μg/mouse, IP route) were administered on day 0. The concentrations of various cytokines and chemokines was determined by multiplex assay. Data represents mean ± SE. * p<0.5, ** p<0.005, *** p<0.0005, **** p<0.00005.

### Intratracheal Infection with a Clinical SA Isolate Producing SEB Elicits Eosinophilic Inflammation

Having established that purified SA-SAgs induce a dose-dependent, SAg-type-dependent pulmonary inflammation, we next examined the effects of intratracheal infection with an SEB-producing clinical SA isolate. HLA-DR3.ENAC mice were infected with a low inoculum of a clinical SA isolate producing SEB (SEB^+^SA) or its isogenic counterpart with the SEB gene inactivated (SEB^-^SA). BAL was collected five days post-infection for analysis. Total BAL cell counts were significantly elevated in mice challenged with SEB^+^SA compared to those challenged with PBS or SEB^-^SA (Fig 10). Among the immune subsets in the BAL, only Eos and immature macrophages/monocytes were significantly higher in mice challenged with SEB^+^SA compared to those challenged with PBS or SEB^-^SA. While PMNs and Ly6G^+^SigF^+^ cells were also elevated in mice challenged with SEB^+^SA, they did not reach statistical significance. Histological examination revealed that lungs from SEB^-^SA challenged mice exhibited only mild perivascular infiltration, whereas lungs from SEB^+^SA challenged mice displayed extensive inflammatory changes with both perivascular and peri-bronchial infiltration. At 40x magnification, large numbers of Eos were readily observable in the infiltrates, reflecting the BAL flow cytometric findings (Fig 10).

**Figure 10.**
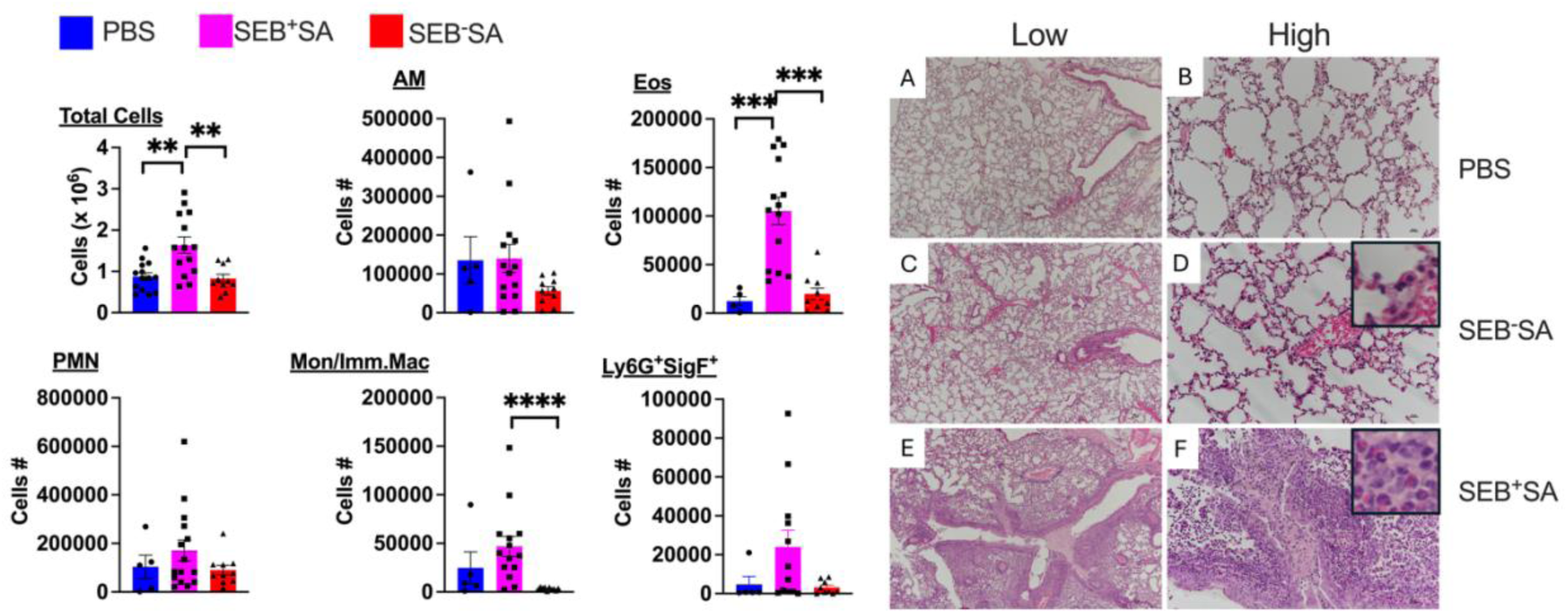
Intratracheal infection with SA elicits airway inflammation in a SAg-dependent manner. HLA-DR3.ENAC mice were challenged intratracheally with PBS, SEB^+^SA or SEB^-^SA and killed 5 days later. Distribution of various BAL leukocyte subsets by flow cytometry. Formalin fixed, paraffin embedded sections were stained with H&E. Data represents mean ± SE. * p<0.5, ** p<0.005, *** p<0.0005, **** p<0.00005.

## DISCUSSION

Despite the remarkable success of CFTR modulator therapies, bacterial infections continue to be a significant driver of chronic airway inflammation and pulmonary exacerbations in pwCF [6, 7, 9, 11, 15, 21]. SA is not only the most prevalent bacterial pathogen capable of persisting for prolonged periods as biofilms in the CF airways, even in those undergoing modulator therapies, it is the only CF pathogen that can produce SA-Ag, which are potent activators of the immune system. Furthermore, clinical CF SA strains frequently harbor multiple SA-SAg genes [30–32], underscoring the potential for recurrent exposure to SAgs in pwCF. Despite this, the contribution of SA-SAgs to the immunopathogenesis of CF lung disease remains largely unexplored. Our study aims to shed light on the overlooked roles of SA-SAgs in the immunopathogenesis of CF lung disease.

First, we confirmed the widespread prevalence of SA-SAg genes in CF SA isolates, encompassing both adult and pediatric populations. Consistent with previous findings, SA-SAgs such as G, I, M, N, and O, encoded by the enterotoxin gene cluster (egc), were frequently identified in all our CF SA isolates [31]. However, other studies have not differentiated between pediatric and adult SA isolates, leaving the distribution of SA-SAgs in these two CF subsets largely unknown. We observed distinct distribution patterns of SA-SAgs between pediatric and adult CF cohorts. For instance, potent SA-SAgs like SEB and SEA were more prevalent in pediatric isolates, which also carried a greater number of SA-SAgs genes compared to adult isolates. A higher prevalence of potent SA-SAgs and a broader distribution of SA-SAg genes in pediatric isolates may explain the stronger association of SA with more severe disease in pediatric CF patients and warrants a deeper investigation [11, 18–20].

Second, our in vitro SA-SAg bioassays with CF clinical isolates grown in ASM as biofilms confirmed the ability of these isolates to produce biologically active SA-SAgs in CF-like settings. Further, our bioassay also suggested that there are intrinsic differences in the extent to which different SA isolates activate the immune system and induce the production of different cytokines such as IFN-γ, which is likely determined by the SAg profile of the isolate (discussed later). Subsequently, we demonstrated that at least SEB, one of the potent SA-SAgs, retained its biological activity in the presence of chemically complex human CF sputum. This is likely applicable for many other SA-SAgs, as several SA-SAgs are known enterotoxins capable of maintaining their biological activities in extremely low gastric pH and in the presence of various proteases and digestive enzymes—conditions likely harsher than those found in CF airways [60]. Overall, a high prevalence of SA-SAgs in CF SA isolates, their ability to produce SA-SAgs in biofilms in vitro under CF-like conditions, the stability of SA-SAgs in CF sputum, and their demonstrated capacity to drive inflammation in other biofilm infection models [61–63], all suggest that SA-SAgs may play a crucial role in the development of lung disease in pwCF. Our in vivo murine studies support this hypothesis.

Recurrent intratracheal exposure to SEB elicited a pulmonary inflammatory response in both B6.ENAC and HLA-DR3.ENAC mice. However, this response was more pronounced in the later. Milder inflammatory responses to SEB or other SA-SAgs in B6 mice have been reported by us and other investigators, attributed to differences in the binding affinities of different SA-SAgs to human versus mouse MHC class II molecules [28, 35, 41–43]. Consequently, B6 mice produce much lower amounts of proinflammatory cytokines such as IFN-γ and TNF-α and exhibit significant resistance to SAg-induced (particularly SEB) acute as well as chronic diseases including toxic shock syndromes [33, 34, 41]. However, this is the first instance where these differences have been demonstrated in a CF model. SA-SAgs could also induce pulmonary inflammation in HLA-DR3.CFGC mice, although not as severe as in HLA-DR3.ENAC mice. This is expected because unlike ENAC mice, CFGC mice do not manifest any spontaneous muco-obstructive pulmonary disease [45]. Hence, HLA-DR3.ENAC mice may be ideally suited for in-depth studies on the role of SA-SAgs in CF lung disease.

The ability of SEB to induce pulmonary inflammation in both ENAC and non-ENAC HLA-DR3 mice suggests that recurrent airway exposure to SA-SAgs could be proinflammatory in all individuals, regardless of CF status. However, since pwCF, particularly children, are more likely to be chronically colonized/infected with SA-SAg-producing SA, this phenomenon is likely to be more relevant in pwCF. It is noteworthy that nasal carriage of SA and recurrent exposure to SA-SAgs have been implicated in the immunopathogenesis of certain non-CF airway inflammatory diseases, including chronic rhinosinusitis. Interestingly, chronic rhinosinusitis is also prevalent among pwCF [64–68]. The following model may outline the mechanisms by which SA-SAgs may contribute to pulmonary inflammation in CF.

Chronic SA infections, including in CF, are often caused by SA growing as biofilms. SA biofilms are localized, inherently slow growing, and metabolically less active. Hence, we have shown that SA biofilms produce small amounts of SA-SAgs locally, albeit continuously. Experiments utilizing fluorescent gene reporter plasmids and other studies have confirmed that SA biofilms readily express SA-SAgs in vitro and in vivo [62, 63, 69–71]. Therefore, in CF patients, the airways (including the sinuses) are likely to be chronically exposed to small quantities of SA-SAgs. As SA-SAgs are resistant to low pH and proteases [60], they are likely to retain their immunostimulatory properties in the CF airways.

Within the airways and sinuses, SA-SAgs can directly activate the epithelial cells to produce IL-8 and other chemokines, driving inflammatory cell recruitment to these sites as shown in other disease models [72–77]. Most importantly, airway epithelial cells (AECs) express MHC class II molecules [78, 79]. Therefore, SA-SAgs could readily activate the mucosal-associated invariant T cells (MAITs), conventional T cells as well as intra epithelial lymphocytes that are present at these mucosal surfaces via HLA class II on AECs, by well-established mechanisms [25–28, 80, 81], initiating a robust inflammatory response at the mucosal surface. SA-SAgs can also be readily absorbed through epithelial cells, including airway epithelial cells, and can even translocate across epithelial cells [82, 83]. Once the SA-SAgs traverse the airway epithelium they can activate adaptive CD4^+^ and CD8^+^ αβ TCR^+^ T cells as well as innate T cells such as Natural Killer T (NKT) cells that are abundantly present in the sub-mucosal areas in the airways [26, 28, 84–86].

Activated innate and adaptive T cells can directly cause cell/tissue injury as well as indirectly by producing several pro-inflammatory type 1, type 2 and type 3 cytokines and chemokines, which can drive the influx of macrophages, neutrophils, eosinophils, and other leukocyte subsets (such as B cells, γδ T cells and NK cells) to the airways to potentiate airway inflammation. A greater lung recruitment of all these cell types in SA-SAg challenged mice supports this model. Even though SA-SAgs do not directly activate B cells, we could readily observe CD19^+^ cells within the lungs. Interestingly, all these cell types, including B cells and NK cells are seen in human CF lungs and are thought to contribute to CF pathogenesis [87, 88]. Similarly, high numbers of γδ T cells and NK1.1^+^ cells in the lungs could be explained by non-specific recruitment of various inflammatory cells. Overall, our in vivo studies clearly supported an important role for SA-SAg in pulmonary inflammation in CF. However, the extent and nature of inflammation induced by SA-SAgs are likely influenced by multiple bacterial and host factors.

SA could produce up to 25 different SA-SAgs [25]. Not all SA strains carry all SAgs genes; some are genomically encoded while others are carried by mobile genetic elements, allowing for the potential loss or gain of SA-SAg genes [89, 90]. The superantigenicity of each SAg varies based on its affinity for specific HLA class II alleles [91–97], and the number of TCR variable regions engaged by a given SAg. For instance, in mice, SEB and SEC activate T cells expressing multiple TCR Vβ, whereas SEA activates only TCR Vβ3. This may explain why SEB (and SEC in CFGC model) are more inflammatory than SEA in mice, a phenomenon that may also apply to humans depending on the SA-SAg, their affinity to HLA class II molecule allotypes, their TCR preference and the frequency of T cells bearing the TCRs [25]. Our in vitro bioassays with different clinical CF isolates also confirm the substantial variation in the abilities of different isolates to activate the immune system and induce the production of different cytokines such as IFN-γ which are likely influenced by the SA-SAg profile.

Furthermore, the concentration of SA-SAgs could critically influence the nature of inflammation. It has been demonstrated that conventional antigens elicit a dose-dependent cytokine response, with low antigen doses favoring a type 2 cytokine response and high doses favoring a type 1 response, particularly in the lungs [98–100]. Similarly, exposure to higher concentrations of SEB could induce the production of higher levels of type 1 cytokines such as IFN-γ, both locally as well as systemically, driving a neutrophilic lung inflammation and increased mortality as shown previously [41, 42, 56]. A dose-dependent increase in the BAL levels of IFN-γ in HLA-DR3.ENAC mice challenged with 10, 100 or 1000 ng of SEB which correlated with the extent of PMN recruitment to the lungs support this hypothesis. Conversely, lower concentrations of SA-SAgs would elicit an eosinophil-predominant type 2 immune response, as demonstrated in a non-CF model of pulmonary inflammation in HLA-DR3 transgenic mice [41].

IFN-γ not only promotes a neutrophilic response [101–103], but also suppresses the production of type 2 cytokines and antagonizes their functions [58, 104–108]. Hence, it has been demonstrated in several models of lung inflammation that neutralization of IFN-γ or IFN-γ-deficient mice mount a more robust type 2 response following antigenic challenge. In a similar manner, even in our CF models, in vivo neutralization of IFN-γ resulted in a more profound type 2 cytokine response, leading to heightened eosinophilic pulmonary inflammation. Differences in the amount of SEB-induced IFN-γ production in vivo could explain why B6.ENAC mice developed a more pronounced eosinophilic inflammation following in vivo IFN-γ neutralization compared to DR3.ENAC mice.

B6 mice produce significantly less IFN-γ than HLA-DR3 mice in response to SEB challenge as discussed above [41]. Consequently, antibodies may be able to completely neutralize IFN-γ, promoting a more robust type 2 cytokine production and an exaggerated eosinophilic inflammation in B6.ENAC mice. In contrast, HLA-DR3 mice produce much higher amounts of IFN-γ following SEB challenge. Hence, the antibodies may not fully neutralize all IFN-γ, allowing residual IFN-γ to promote a neutrophilic response and only modestly suppressing type 2 cytokines [101–103]. A similar dose-dependent differential induction of type 1 versus type 2 responses could explain a skewed eosinophilic lung inflammation following in vivo infection with SA in our model. We have previously demonstrated that intratracheal infection of non-CF HLA-DR3 transgenic mice with a higher inoculum of SEB-producing SA rapidly induced the production of high levels of IFN-γ which was associated with high mortality, which could be reversed by in vivo neutralization of SEB [109, 110]. Likewise, infection with a lower inoculum of SEB-producing SA in this study could have resulted in lower IFN-γ production and a more pronounced type 2 response, similar to the low-dose SEB.

In our CF models, we consistently observed a significantly higher numbers of Sig F^+^ Ly6G^+^ cells within the granulocyte gate. Sig F^+^ Ly6G^+^ (or Sig F^+^ Gr-1^+^) are shown to be eosinophils and have been demonstrated in mouse models of asthma and other type 2-dominant diseases [111–114]. A significant increase in the numbers of Sig F^+^ Ly6G^+^ cells in our model, that are traditionally associated with strong type 2 responses, underscores the ability of SA-SAgs to drive type 2-type inflammation in the CF lungs, more so when IFN-γ is blocked. However, additional studies are needed to fully characterize this unique population of cells and their role in CF.

It is well established that some pwCF exhibit a skewed type 2 response, and patients with eosinophilic inflammation tend to have poorer outcomes [115–120]. However, the underlying mechanisms have remained unclear and have often been attributed to infections with Aspergillus [121]. Our studies suggest that SA, through SA-SAgs could also drive eosinophilic lung disease. Further, IgE antibodies to SA-SAgs have been implicated in the development of type 2 inflammation in certain non-CF lung diseases [122, 123]. A similar mechanism could be involved in the development of type 2 inflammation in the CF lungs. Overall, a more comprehensive prospective clinical study is warranted to establish the role of SA-SAgs in the pathogenesis of CF lung disease. While the lack of sensitive assays to detect SA-SAgs in biological samples, particularly in chemically complex materials such as CF sputum, poses a significant limitation, sputum transcriptomic studies may provide valuable insights.

In summary, we have demonstrated that airway exposure to SA-SAgs can trigger pulmonary inflammation, the nature of which is likely influenced by various host and bacterial factors. As colonization or infection with SA can be either persistent or intermittent in CF airways [124], with the strains involved potentially changing over time, there is expected to be variabilities in the SA-SAg produced, hence the nature of inflammatory response. Further, some SA strains may remain confined to the sinuses and never reach the lower airways posing additional heterogeneity [125]. Additionally, we have observed metabolic alterations in SA isolated over time in a subset of CF patients who developed SA-induced pneumonia or bacteremia [126]. Increase in the production of SA-SAgs due to SA metabolic changes may explain the worsening of clinical presentations with systemic involvements in these patients. These observations may also help explain the variability in the presentation of CF lung disease, despite the widespread prevalence of SA among pwCF. Nevertheless, our novel findings underscore the necessity for more extensive studies to fully elucidate the role of SA and SA-SAgs in the development of lung disease in pwCF.

## ACKNOWLEDGEMENTS

The project is funded Research Pilot and Feasibility Grant from Cystic Fibrosis Foundation to GR. We thank the Center for Phage Biology and Therapy at Yale for providing us the adult CF SA isolates. We thank Yale Flow Cytometry for their assistance with flowcytometry service. The Core is supported in part by an NCI Cancer Center Support Grant # NIH P30 CA016359. The BD Symphony was funded by shared instrument grant # NIH S10 OD026996. We thank the Yale Pathology Tissue Services Core facility for help with tissue sectioning and immunostaining.

## AUTHOR CONTRIBUTIONS

YS, BH, SC, JT, NS, JN - Conducting experiments and acquiring data.

ZH, GS - Analyzing data and writing the manuscript.

CB, PP - Providing reagents, analyzing data and writing the manuscript.

SS, RA, JA - Providing reagents (developing, characterizing and providing MNHOCH and MNHOCHΔ SEB and writing the manuscript.

XZ - Analyzing data and writing the manuscript.

JK - Development of hypothesis, analyzing data and writing the manuscript.

GR – Development of hypothesis, designing research studies, conducting experiments, acquiring data, analyzing data and writing the manuscript.

## ABBREVIATIONS

AEC: Airway epithelial cells
ASM: Artificial sputum medium
CF: cystic fibrosis (CF)
CFGC: cystic fibrosis gut corrected
CFTR: Cystic Fibrosis Transmembrane Conductance Regulator
ENAC: Epithelial Sodium Channel
MAITs: Mucosal associated invariant T cells
pwCF: people with CF
SA: Staphylococcus aureus
SA-SAg: Staphylococcus aureus superantigens
SAg: Superantigens
SEA: Staphylococcal enterotoxin A
SEB: Staphylococcal enterotoxin B
SEC: Staphylococcal enterotoxin C
TST: Toxic shock syndrome toxin-1

