## Supplemental Data for "Prevalence, Production and the Role of *Staphylococcus aureus* Superantigens in Cystic Fibrosis Lung Disease"

### Supplementary Figure Legends

Supplementary Figure 1. WB showing the absence of SEB protein in the MNHOCH Δ SEB clone. MNHOCH WT and MNHOCHΔSEB were cultured overnight in 7 mL of TSB media at 37°C with constant shaking at 230 rpm. Next day, $OD_{600}$ was measured, and adjusted to an $OD_{600}$ of 6 by adding TSB media. The bacterial culture supernatants (BCS) were collected by centrifugation at 3500 rpm for 10 minutes at room temperature and filtered through a 0.2 μm filter. BCS was concentrated using a 3K MWCO filter and heated at 70°C for 10 minutes under reducing conditions. Immunoblotting was performed with 1 μg/mL anti-SEB antibody (IDP-02), incubated overnight at RT. Peroxidase activity was detected using goat anti-human HRP secondary antibody with ECL substrate. Lane 1: MW ladder (kDa), Lane 2: MNHOCH WT expressing SEB (~28 kDa), Lane 3: MNHOCH Δ SEB (absence of SEB), Lane 4: purified 25 ng SEB protein (control). SEB is not detected in the MNHOCH Δ SEB clone. The primary antibody cross-reacts with Protein A (~50 kDa).

Supplementary Figure 2. Gating strategy for phenotyping different leukocyte subjects in BAL and lung digests by spectral flow cytometry. (A) Myeloid cell gating strategy (B) Lymphoid cell gating strategy.

Supplementary Figure 3. SA-SAg retains its superantigenicity in the presence of CF sputum extracts. Purified SEB (10 μg) was added to 1 ml of CF sputum extracts or PBS and incubated at 37°C. (A) IL-2 production by splenocytes from HLA-DR3 transgenic mice following stimulation with SEB (1 μg/ml) that had been previously incubated with CF sputum extract or PBS. (B) HLA-DR3 transgenic mice were challenged with SEB that had been incubated with CF sputum extract or PBS as above (@ 10 μg/mouse). Three days later, spleens were collected from mice and the distribution of TCR Vβ6$^{+}$ and TCR Vβ8$^{+}$ T cells within the CD4$^{+}$ or CD8$^{+}$ T cell subsets were determined by flow cytometry

Supplementary Figure 4. Phenotypic differences between BAL cells collected from HLA-DR3.ENAC challenged with SEB or PBS. HLA-DR3.ENAC mice were challenged intratracheally with PBS or 100 ng of SEB on days 0, 3 and 6 and killed on day 8. BAL cells collected on day 8 were analyzed by spectral flowcytometry. Representative dot plots and histograms are shown.

Supplementary Figure 5. Lung infiltration of CD3$^{+}$ cells, CD19$^{+}$ cells and eosinophils following intratracheal SEB challenge. HLA-DR3.ENAC or B6.ENAC mice were challenged intratracheally with PBS or 100 ng of SEB on days 0, 3 and 6 and killed on day 8. After collecting BAL, lungs were perfused, the right lobes were formalin fixed, sectioned and stained with anti-CD3 antibodies, anti-CD19 antibodies or anti-MBP antibody. Representative photomicrographs are shown.

Supplementary Figure 6. Differential inflammatory potentials of SEA and SEB. HLA-DR3.ENAC mice were challenged intratracheally with PBS or 100 ng of SEA or SEB on days 0, 3 and 6 and killed on day 8. Total cell counts determined using a Coulter counter. Distribution of various BAL leukocyte subsets as determined by spectral flow cytometry. Data represents mean ± SE. * $p<0.5$, ** $p<0.005$, *** $p<0.0005$, **** $p<0.00005$. Pathological changes in the lungs were also studied. Representative images at different magnifications are show.

Supplementary Figure 7. SA-SAg induced pulmonary inflammation in HLA-DR3.CFGC mice. HLA-DR3.CFGC mice were challenged intratracheally with 100 ng of SEA, SEB or SEC on days 0, 3 and 6 and killed on day 8. BAL leukocyte subsets were determined by spectral flow cytometry. Data represents mean ± SE. * $p<0.5$, ** $p<0.005$ compared to PBS group.

Supplementary Figure 8. Dose-dependent inflammatory changes in the lungs following intratracheal challenge with SEB. HLA-DR3.ENAC mice were challenged intratracheally with 10 or 1000 ng of SEB on days 0, 2 and 4 and killed on day 6. After collecting BAL, lungs were perfused, the right lobes were formalin fixed, sectioned and stained with hematoxylin and eosin. Representative photomicrographs are shown.

Supplementary File 1. SA-SAg sequences used to identify SA-SAg gene distribution in CF SA isolates.

SF 1

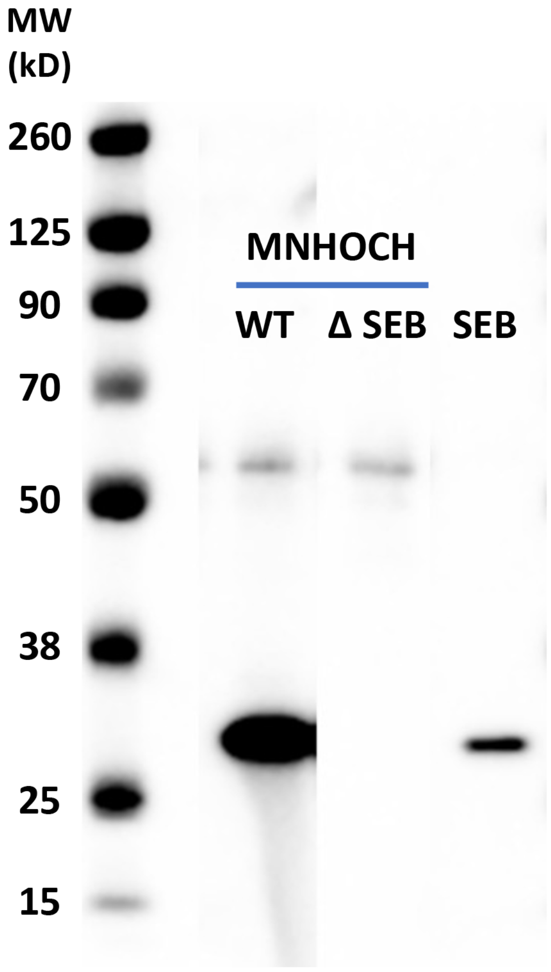

SF 2A

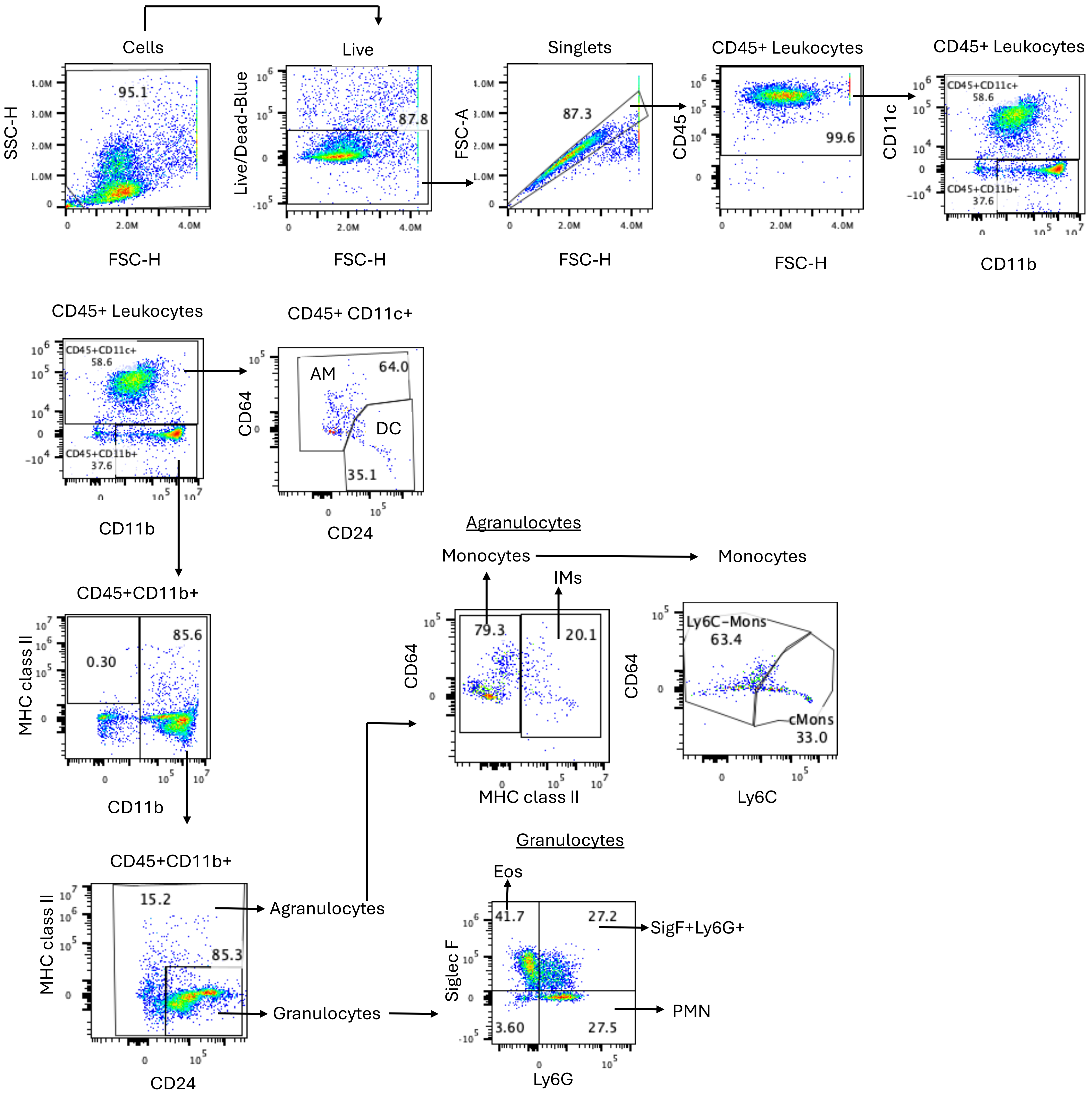

Cells
95.1
SSC-H
FSC-H
Live
87.8
Live/Dead-Blue
FSC-H
Singlets
87.3
FSC-A
FSC-H
CD45+ Leukocytes
99.6
CD45
FSC-H
CD45+ Leukocytes
CD45+CD11c+
58.6
CD45+CD11b+
37.6
CD11c
CD11b
CD45+ Leukocytes
CD45+CD11c+
58.6
CD45+CD11b+
37.6
CD11b
CD45+ CD11c+
AM
64.0
DC
35.1
CD64
CD24
CD45+CD11b+
0.30
85.6
MHC class II
CD11b
CD45+CD11b+
15.2
85.3
Agranulocytes
Granulocytes
MHC class II
CD24
Agranulocytes
Monocytes
IMs
Monocytes
79.3
20.1
CD64
MHC class II
Ly6C−Mons
63.4
cMons
33.0
CD64
Ly6C
Granulocytes
Eos
41.7
27.2
SigF+Ly6G+
3.60
27.5
PMN
Siglec F
Ly6G

SF 2B

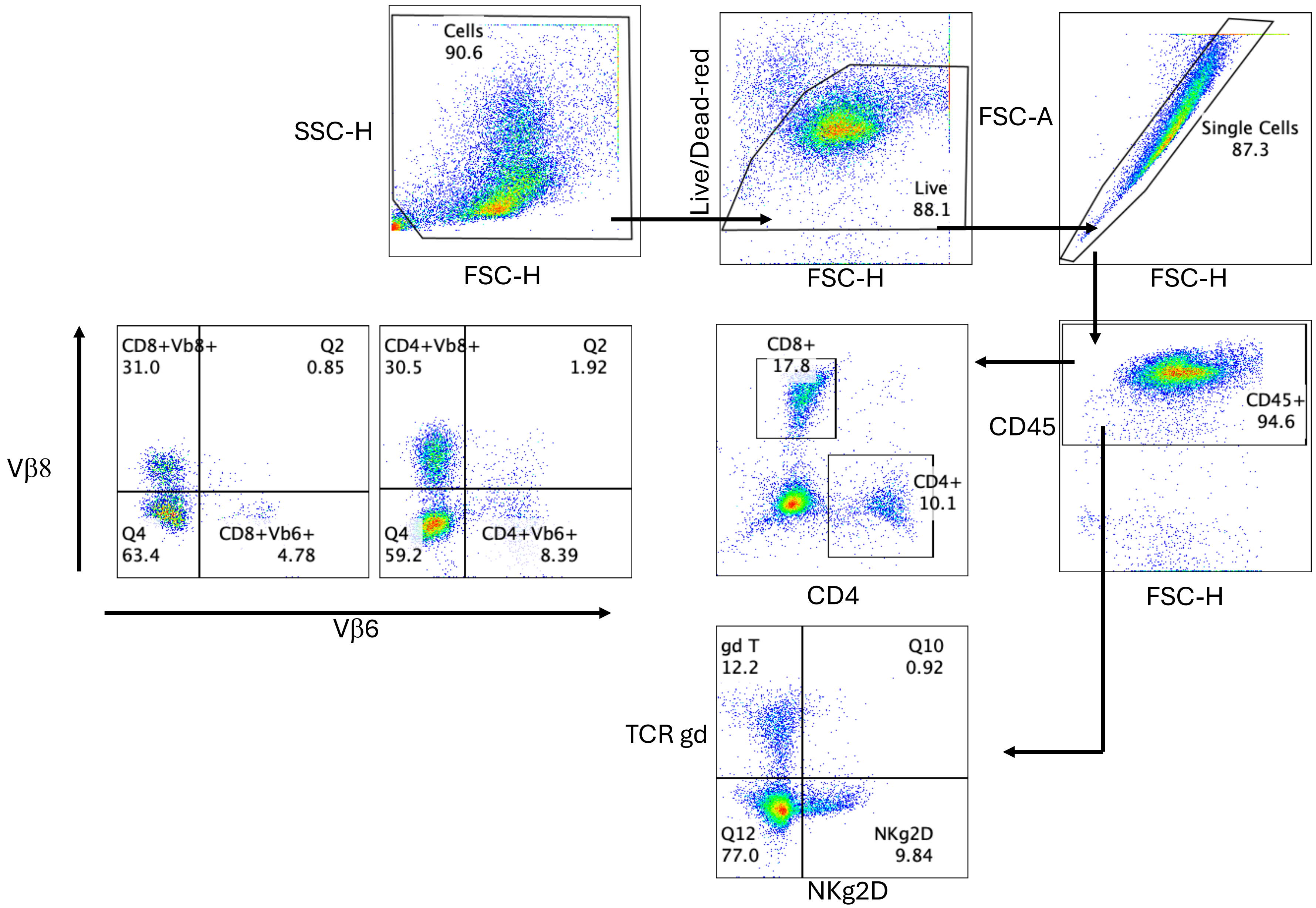

Cells
90.6
SSC-H
FSC-H
Live/Dead-red
Live
88.1
FSC-H
FSC-A
Single Cells
87.3
FSC-H
CD45
CD45+
94.6
FSC-H
CD8+
17.8
CD4+
10.1
CD4
CD8+Vb8+
31.0
Q2
0.85
Q4
63.4
CD8+Vb6+
4.78
CD4+Vb8+
30.5
Q2
1.92
Q4
59.2
CD4+Vb6+
8.39
Vβ8
Vβ6
TCR gd
gd T
12.2
Q10
0.92
Q12
77.0
NKg2D
9.84
NKg2D

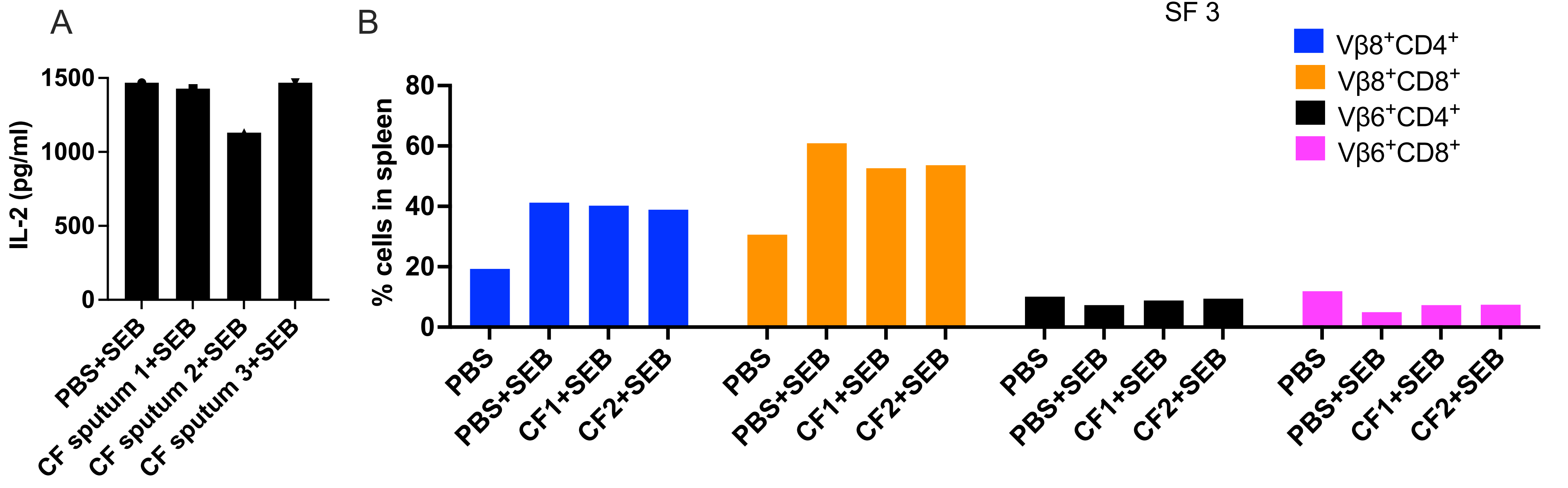

A
IL-2 (pg/ml)
1500
1000
500
0
PBS+SEB
CF sputum 1+SEB
CF sputum 2+SEB
CF sputum 3+SEB
B
SF 3
% cells in spleen
80
60
40
20
0
Vβ8+CD4+
Vβ8+CD8+
Vβ6+CD4+
Vβ6+CD8+
PBS
PBS+SEB
CF1+SEB
CF2+SEB

SF 4

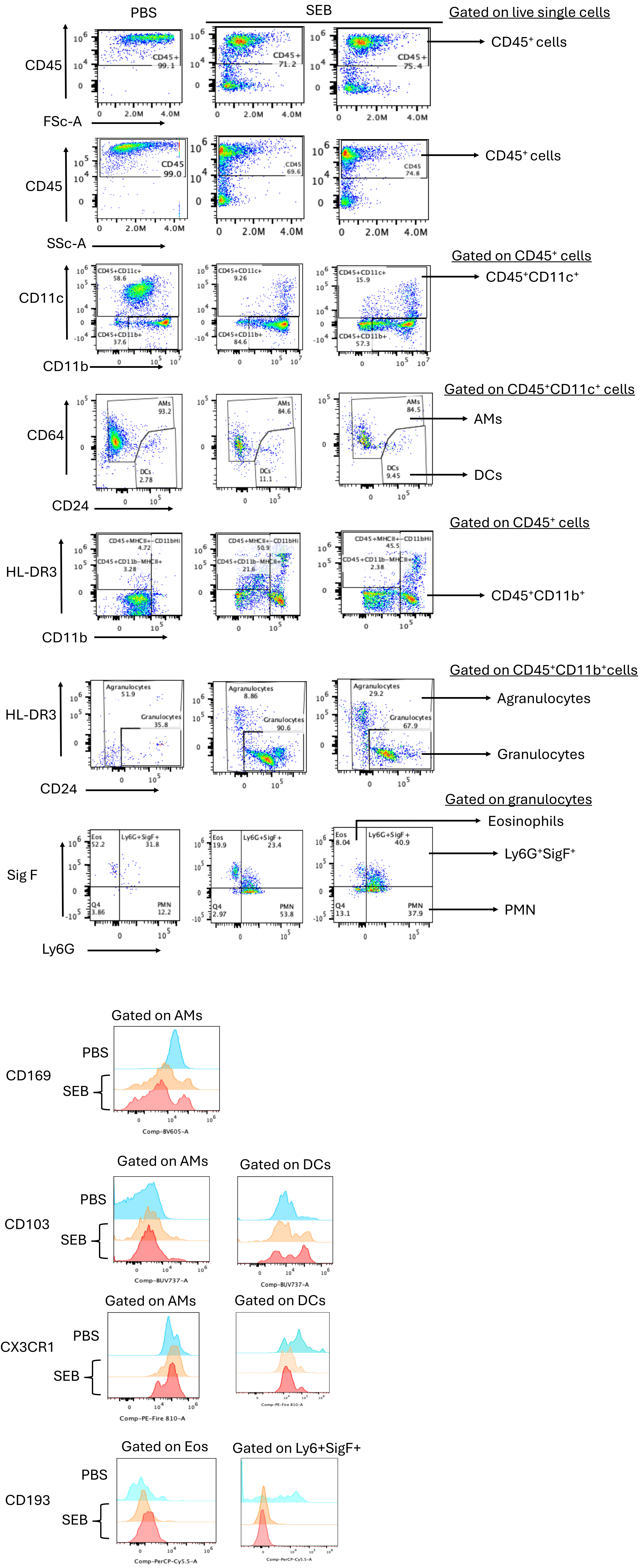
PBS
SEB
Gated on live single cells
CD45
FSc-A
CD45+ 99.1
CD45+ 71.2
CD45+ 75.4
CD45⁺ cells
SSc-A
CD45 99.0
CD45 69.6
CD45 74.8
CD45⁺ cells
Gated on CD45⁺ cells
CD11c
CD11b
CD45+CD11c+ 58.6
CD45+CD11b+ 37.6
CD45+CD11c+ 9.26
CD45+CD11b+ 84.6
CD45+CD11c+ 15.9
CD45+CD11b+ 57.3
CD45⁺CD11c⁺
Gated on CD45⁺CD11c⁺ cells
CD64
CD24
AMs 93.2
DCs 2.78
AMs 84.6
DCs 11.1
AMs 84.5
DCs 9.45
AMs
DCs
Gated on CD45⁺ cells
HL-DR3
CD11b
CD45+MHCII+-CD11bHi 4.72
CD45+CD11b-MHCII+ 3.28
CD45+MHCII+-CD11bHi 50.9
CD45+CD11b-MHCII+ 21.6
CD45+MHCII+-CD11bHi 45.5
CD45+CD11b-MHCII+ 2.38
CD45⁺CD11b⁺
Gated on CD45⁺CD11b⁺cells
HL-DR3
CD24
Agranulocytes 51.9
Granulocytes 35.8
Agranulocytes 8.86
Granulocytes 90.6
Agranulocytes 29.2
Granulocytes 67.9
Agranulocytes
Granulocytes
Gated on granulocytes
Sig F
Ly6G
Eos 52.2
Ly6G+SigF+ 31.8
Q4 3.86
PMN 12.2
Eos 19.9
Ly6G+SigF+ 23.4
Q4 2.97
PMN 53.8
Eos 8.04
Ly6G+SigF+ 40.9
Q4 13.1
PMN 37.9
Eosinophils
Ly6G⁺SigF⁺
PMN
Gated on AMs
CD169
PBS
SEB
Comp-BV605-A
Gated on AMs
Gated on DCs
CD103
PBS
SEB
Comp-BUV737-A
Comp-BUV737-A
Gated on AMs
Gated on DCs
CX3CR1
PBS
SEB
Comp-PE-Fire 810-A
Comp-PE-Fire 810-A
Gated on Eos
Gated on Ly6+SigF+
CD193
PBS
SEB
Comp-PerCP-Cy5.5-A
Comp-PerCP-Cy5.5-A

SF 5

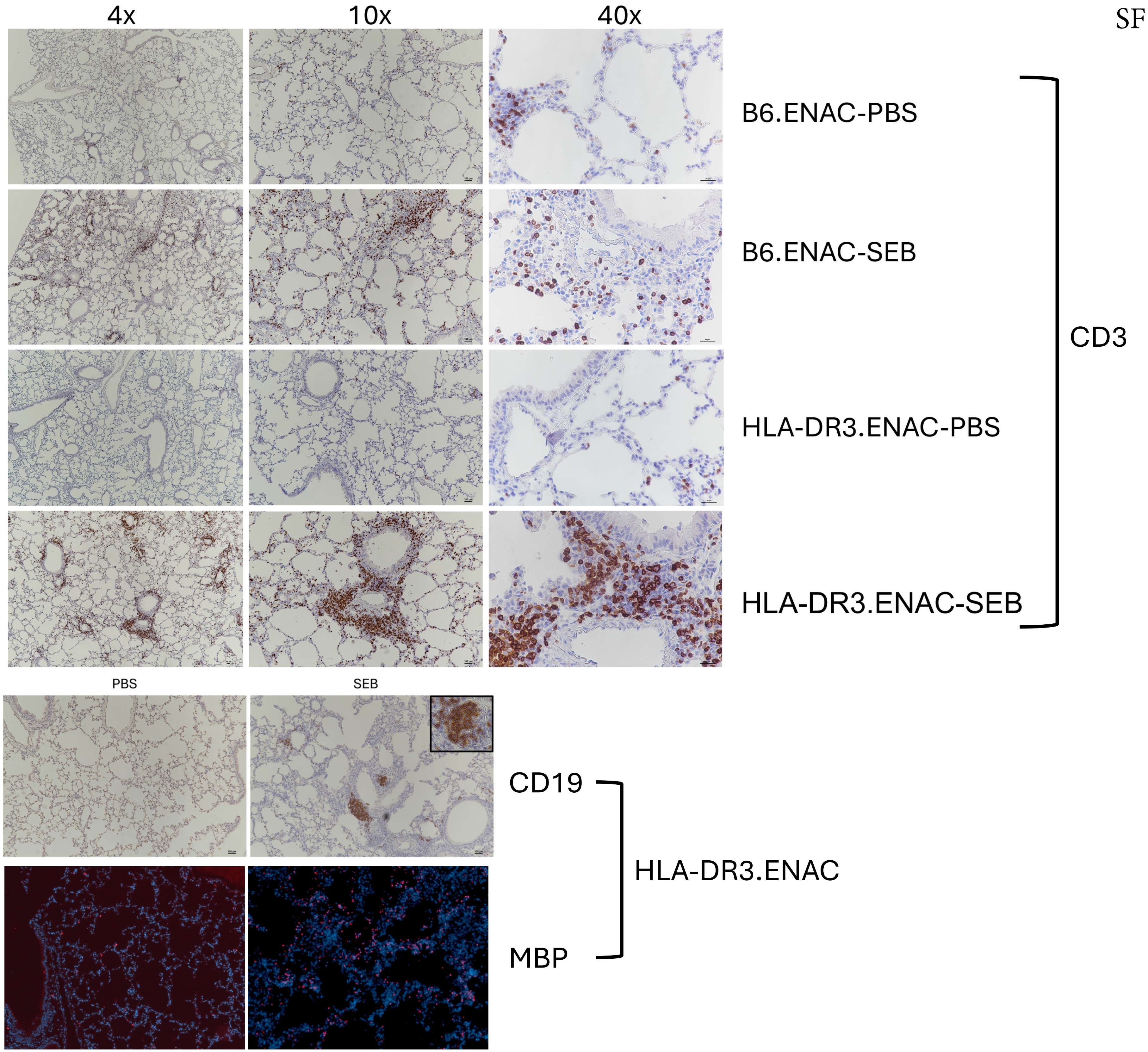
4x
10x
40x
B6.ENAC-PBS
B6.ENAC-SEB
HLA-DR3.ENAC-PBS
HLA-DR3.ENAC-SEB
CD3
PBS
SEB
CD19
HLA-DR3.ENAC
MBP

SF 6

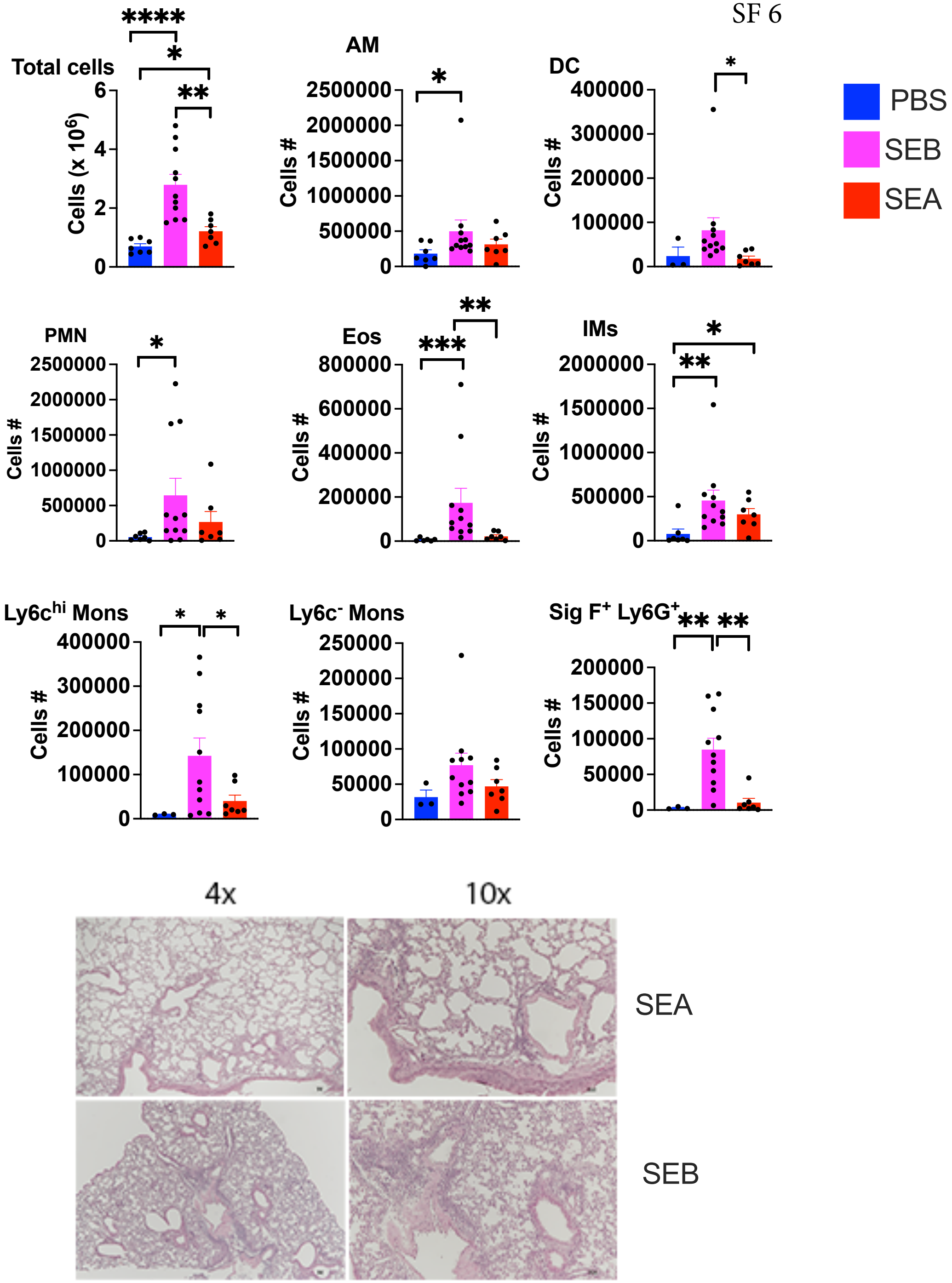

PBS
SEB
SEA
Total cells
Cells (x 10^6)
AM
Cells #
DC
PMN
Eos
IMs
Ly6c^hi Mons
Ly6c^- Mons
Sig F^+ Ly6G^+
4x
10x
SEA
SEB

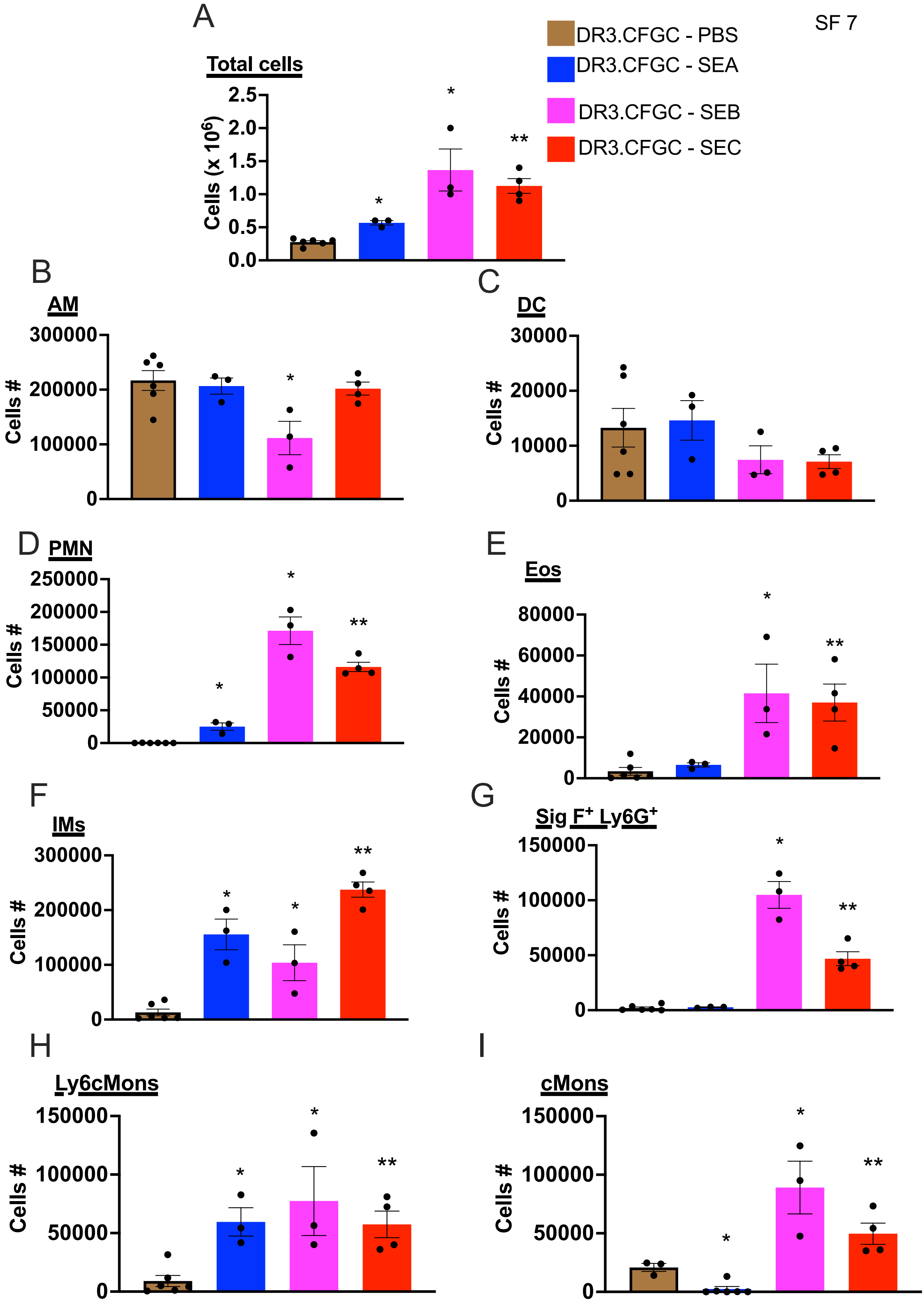
SF 7
A
Total cells
Cells (x 10⁶)
2.5
2.0
1.5
1.0
0.5
0.0
*
*
**
DR3.CFGC - PBS
DR3.CFGC - SEA
DR3.CFGC - SEB
DR3.CFGC - SEC
B
AM
300000
200000
100000
0
Cells #
C
DC
30000
20000
10000
0
D
PMN
250000
200000
150000
100000
50000
E
Eos
80000
60000
40000
20000
F
IMs
G
Sig F⁺ Ly6G⁺
150000
100000
50000
H
Ly6cMons
I
cMons

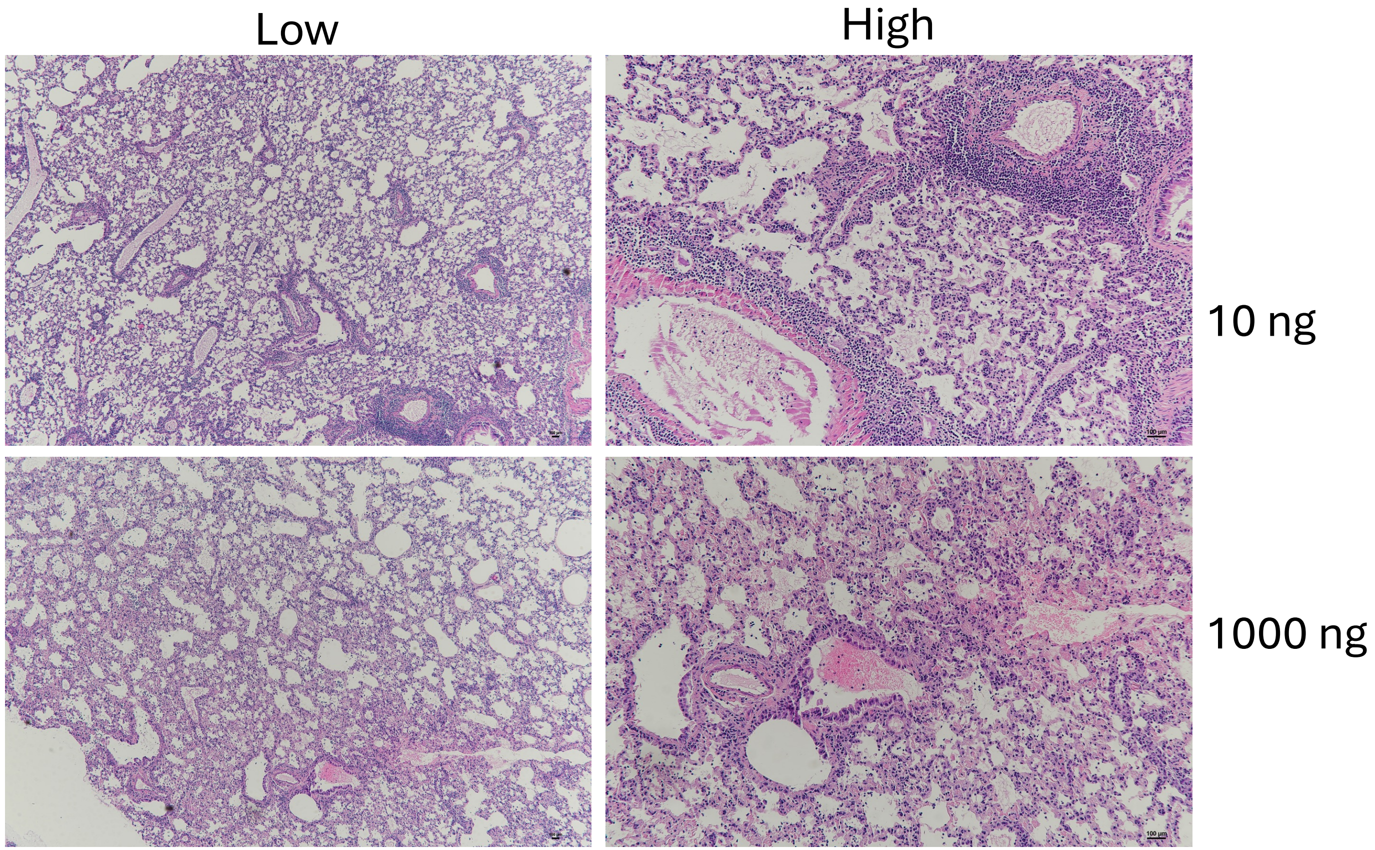
Low
High
10 ng
1000 ng

>vfdb~~~tsst-1~~~NP_375120 (tsst-1) toxic shock syndrome toxin-1 [TSST-1 (VF0022)] [Staphylococcus aureus subsp. aureus N315]
ATGAATAAAAAATTACTAATGAATTTTTTTATCGTAAGCCCTTTGTTGCTTGCGACAATC
GCTACAGATTTTACCCCTGTTCCCTTATCATCTAATCAAATAATCAAAACTGCAAAAGCA
TCTACAAACGATAATATAAAGGATTTGCTAGACTGGTATAGTAGTGGGTCTGACACTTTT
ACAAATAGTGAAGTTTTAGATAATTCCTTAGGATCTATGCGTATAAAAAACACAGATGGC
AGCATCAGCCTTATAATTTTTCCGAGTCCTTATTATAGCCCTGCTTTTACAAAAGGGGAA
AAAGTTGACTTAAACACAAAAAGAACTAAAAAAAGCCAACATACTAGCGAAGGAACTTAT
ATCCATTTCCAAATAAGTGGCGTTACAAATACTGAAAAATTACCTACTCCAATAGAACTA
CCTTTAAAAGTTAAGGTTCATGGTAAAGATAGCCCCTTAAAGTATTGGCCAAAGTTCGAT
AAAAAACAATTAGCTATATCAACTTTAGACTTTGAAATTCGTCATCAGCTAACTCAAATA
CATGGATTATATCGTTCAAGCGATAAAACGGGTGGTTATTGGAAAATAACAATGAATGAC
GGATCCACATATCAAAGTGATTTATCTAAAAAGTTTGAATACAATACTGAAAAACCACCT
ATAAATATTGATGAAATAAAAACTATAGAAGCAGAAATTAATTAA
>vfdb~~~sea~~~NP_646706 (sea) staphylococcal enterotoxin A precursor [SE (VF0020)] [Staphylococcus aureus subsp. aureus MW2]
ATGAAAAAAACAGCATTTACATTACTTTTATTCATTGCCCTAACGTTGACAACAAGTCCA
CTTGTAAATGGTAGCGAGAAAAGCGAAGAAATAAATGAAAAAGATTTGCGAAAAAAGTCT
GAATTGCAGGGAACAGCTTTAGGCAATCTTAAACAAATCTATTATTACAATGAAAAAGCT
AAAACTGAAAATAAAGAGAGTCACGATCAATTTTTACAGCATACTATATTGTTTAAAGGC
TTTTTTACAGATCATTCGTGGTATAACGATTTATTAGTAGATTTTGATTCAAAGGATATT
GTTGATAAATATAAAGGGAAAAAAGTAGACTTGTATGGTGCTTATTATGGTTATCAATGT
GCGGGTGGTACACCAAACAAAACAGCTTGTATGTATGGTGGTGTAACGTTACATGATAAT
AATCGATTGACCGAAGAGAAAAAAGTGCCGATCAATTTATGGCTAGACGGTAAACAAAAT
ACAGTACCTTTGGAAACGGTTAAAACGAATAAGAAAAATGTAACTGTTCAGGAGTTGGAT
CTTCAAGCAAGACGTTATTTACAGGAAAAATATAATTTATATAACTCTGATGTTTTTGAT
GGGAAGGTTCAGAGGGGATTAATCGTGTTTCATACTTCTACAGAACCTTCGGTTAATTAC
GATTTATTTGGTGCTCAAGGACAGTATTCAAATACACTATTAAGAATATATAGAGATAAT
AAAACGATTAACTCTGAAAACATGCATATTGATATATATTTATATACAAGTTAA
>vfdb~~~seb~~~AAA88550 (seb) staphylococcal enterotoxin B [SE (VF0020)] [Staphylococcus aureus S6]
ATGTATAAGAGATTATTTATTTCACATGTAATTTTGATATTCGCACTGATATTAGTTATT
TCTACACCCAACGTTTTAGCAGAGAGTCAACCAGATCCTAAACCAGATGAGTTGCACAAA
TCGAGTAAATTCACTGGTTTGATGGAAAATATGAAAGTTTTGTATGATGATAATCATGTA
TCAGCAATAAACGTTAAATCTATAGATCAATTTCTATACTTTGACTTAATATATTCTATT
AAGGACACTAAGTTAGGGAATTATGATAATGTTCGAGTCGAATTTAAAAACAAAGATTTA
GCTGATAAATACAAAGATAAATACGTAGATGTGTTTGGAGCTAATTATTATTATCAATGT
TATTTTTCTAAAAAAACGAATGATATTAATTCGCATCAAACTGACAAACGAAAAACTTGT
ATGTATGGTGGTGTAACTGAGCATAATGGAAACCAATTAGATAAATATAGAAGTATTACT
GTTCGGGTATTTGAAGATGGTAAAAATTTATTATCTTTTGACGTACAAACTAATAAGAAA
AAGGTGACTGCTCAAGAATTAGATTACCTAACTCGTCACTATTTGGTGAAAAATAAAAAA
CTCTATGAATTTAACAACTCGCCTTATGAAACGGGATATATTAAATTTATAGAAAATGAG
AATAGCTTTTGGTATGACATGATGCCTGCACCAGGAGATAAATTTGACCAATCTAAATAT
TTAATGATGTACAATGACAATAAAATGGTTGATTCTAAAGATGTGAAGATTGAAGTTTAT
CTTACGACAAAGAAAAAGTGA
>vfdb~~~sec~~~NP_645576 (sec) staphylococcal enterotoxin C precursor [Enterotoxin C (CVF057)] [Staphylococcus aureus subsp. aureus MW2]
ATGAATAAGAGTCGATTTATTTCATGCGTAATTTTGATATTCGCACTTATACTAGTTCTT
TTTACACCCAACGTATTAGCAGAGAGCCAACCAGACCCTACGCCAGATGAGTTGCACAAA
TCAAGTGAGTTTACTGGTACGATGGGTAATATGAAATATTTATATGATGATCATTATGTA

TCAGCAACTAAAGTTAAGTCTGTAGATAAATTTTTGGCACATGATTTAATTTATAACATT
AGTGATAAAAAACTAAAAAATTATGACAAAGTGAAAACAGAGTTATTAAATGAAGATTTA
GCAAAAAAGTACAAAGATGAAGTAGTTGATGTGTATGGATCAAATTACTATGTAAACTGC
TATTTTTCATCCAAAGATAATGTAGGTAAAGTTACAGGTGGTAAAACTTGTATGTATGGA
GGAATAACAAAACATGAAGGAAACCACTTTGATAATGGGAACTTACAAAATGTACTTATA
AGAGTTTATGAAAATAAAAGAAACACAATTTCTTTTGAAGTGCAAACTGATAAGAAAAGT
GTAACAGCTCAAGAACTAGACATAAAAGCTAGGAATTTTTTAATTAATAAAAAAAATTTG
TATGAGTTTAACAGTTCACCATATGAAACAGGATATATAAAATTTATTGAAAATAACGGC
AATACTTTTTGGTATGATATGATGCCTGCACCAGGCGATAAGTTTGACCAATCTAAATAT
TTAATGATGTACAACGACAATAAAACGGTTGATTCTAAAAGAGTGAAGATAGAAGTCCAC
CTTACAACAAAGAATGGATAA
>vfdb~~~sed~~~AAB06195 (sed) staphylococcal enterotoxin D precursor
[SE (VF0020)] [Staphylococcus aureus RN4220]
ATGAAAAAATTTAACATTCTTATTGCATTACTCTTTTTTACTAGTTTGGTAATATCTCCT
TTAAACGTTAAAGCCAATGAAAACATTGATTCAGTAAAAGAGAAAGAATTGCATAAAAAA
TCTGAATTAAGTAGTACCGCGCTAAATAATATGAAACATTCTTATGCAGATAAAAATCCA
ATAATAGGAGAAAATAAAAGTACAGGAGATCAATTTTTAGAAAATACTTTGCTTTACAAA
AAATTTTTTACTGACCTTATCAATTTTGAAGATTTATTAATAAACTTCAATTCAAAAGAA
ATGGCTCAACATTTCAAATCTAAAAATGTAGATGTTTACCCTATAAGATATAGCATTAAT
TGTTATGGTGGTGAAATAGATAGGACTGCTTGTACATATGGAGGTGTCACTCCACACGAA
GGTAATAAATTAAAAGAACGAAAAAAAATACCAATCAATTTGTGGATAAATGGTGTACAA
AAAGAAGTTTCTTTAGATAAAGTTCAAACAGATAAAAAAAATGTTACCGTACAAGAATTA
GATGCACAAGCAAGGCGCTATTTGCAAAAGGATTTAAAATTGTATAATAATGATACTCTC
GGAGGAAAAATACAGCGCGGAAAAATAGAGTTTGATTCTTCTGATGGGTCTAAAGTCTCT
TATGATTTATTTGATGTTAAGGGTGATTTTCCCGAAAAACAATTACGAATATACAGTGAT
AATAAAACATTATCCACAGAGCACCTTCATATTGACATCTATTTATATGAAAAGTAG
>vfdb~~~see~~~AAA26617 (see) staphylococcal enterotoxin E precursor
[SE (VF0020)] [Staphylococcus aureus]
ATGAAAAAAACAGCATTTATACTACTTTTATTCATTGCCCTAACGTTGACAACAAGTCCA
CTTGTAAATGGTAGCGAGAAAAGCGAAGAAATAAATGAAAAAGATTTGCGAAAAAAGTCT
GAATTACAAAGAAATGCTTTAAGCAATCTTAGGCAAATTTATTATTATAATGAAAAAGCT
ATAACTGAAAACAAAGAGAGTGATGATCAGTTTTTAGAGAATACTTTGTTATTTAAAGGT
TTTTTCACAGGTCATCCATGGTATAACGATTTATTAGTAGATCTTGGTTCAAAAGATGCT
ACTAATAAATATAAAGGGAAAAAAGTAGACTTATATGGTGCTTATTATGGATATCAATGT
GCTGGAGGCACACCAAATAAAACAGCATGTATGTACGGGGGTGTAACATTACATGATAAT
AACCGATTGACCGAAGAAAAAAAAGTACCAATTAACTTGTGGATAGACGGAAAACAAACT
ACAGTACCTATAGATAAAGTTAAAACAAGCAAAAAAGAAGTAACTGTTCAAGAGCTAGAT
CTTCAGGCAAGGCATTATTTACACGGAAAATTTGGTTTATATAACTCAGACAGCTTTGGC
GGTAAGGTGCAAAGAGGCTTGATTGTGTTTCATTCTTCTGAAGGGTCCACGGTAAGTTAT
GATTTGTTTGATGCTCAAGGGCAATATCCAGATACATTATTAAGAATTTACAGAGATAAT
AAAACTATTAATTCAGAGAACCTCCATATTGATTTGTATTTATACACAACTTGA
>vfdb~~~seh~~~NP_644866 (seh) staphylococcal enterotoxin H precursor
[SE (VF0020)] [Staphylococcus aureus subsp. aureus MW2]
ATGATTAATAAAATTAAAATATTATTTTCGTTTTTAGCATTATTACTTTCATTCACATCA
TATGCGAAAGCAGAAGATTTACACGATAAAAGTGAGTTAACAGATTTAGCTTTAGCTAAT
GCATATGGTCAATATAATCACCCATTCATTAAAGAAAATATTAAGAGTGATGAAATAAGT
GGAGAAAAAGATTTAATATTTAGAAATCAAGGTGATAGTGGCAATGATTTGAGAGTAAAG
TTTGCAACTGCTGATTTAGCTCAGAAGTTTAAAAATAAAAATGTAGATATATATGGGGCA
TCTTTTTATTATAAGTGTGAAAAAATAAGTGAAAATATTTCTGAATGTCTATATGGAGGT
ACAACACTAAATAGTGAAAAATTGGCACAGGAAAGGGTGATTGGTGCTAATGTTTGGGTA

GATGGTATTCAAAAAGAAACAGAATTAATACGAACAAATAAGAAAAATGTGACATTGCAA
GAATTAGATATAAAGATCAGAAAAATATTGTCCGATAAATATAAAATTTATTATAAAGAC
AGCGAAATAAGTAAAGGTCTAATTGAATTTGATATGAAAACTCCTAGAGATTACTCATTC
GACATTTATGATTTAAAAGGAGAAAATGACTATGAGATAGATAAAATTTATGAAGACAAT
AAAACTTTAAAATCTGATGATATAAGTCATATTGATGTAAATCTATATACTAAGAAAAAA
GTATAA
>vfdb~~~selk~~~NP_646755 (selk) staphylococcal enterotoxin K precursor
[SE (VF0020)] [Staphylococcus aureus subsp. aureus MW2]
ATGAAAAAATTAATAAGCATCTTATTAATAAATATAATAATTTTAGGTGTCTCTAATAAT
GCCAGCGCTCAAGGTGATATAGGAATTGATAATCTCAGGAATTTTTATACAAAAAAAGAC
TTTATAAATTTAAAAGATGTAAAAGACAATGATACTCCTATAGCTAATCAACTACAATTT
TCAAATGAATCCTATGATTTAATTTCAGAATCAAAAGATTTTAATAAATTTAGTAATTTC
AAAGGAAAAAAACTTGATGTTTTTGGTATTAGTTATAATGGCCAGTGTAACACTAAATAC
ATATATGGTGGAATCACAGCTACTAACGAATATCTAGATAAACCTAGAAATATACCTATA
AATATATGGATCAATGGAAATCACAAAACTATTTCTACCAATAAAGTTTCGACAAATAAA
AAATTTGTTACCGCTCAAGAGATTGATATCAAATTAAGAAGGTACCTTCAAGAAGAATAC
AACATTTATGGACATAACGGCACTAAAAAAGGAGAAGAATACGGTCATAAATCAAAATTT
TATTCTGGGTTTAATATTGGTAAAGTAACGTTTCATTTAAATAATAATGACACCTTTTCA
TATGATTTATTCTACACAGGAGATGATGGGCTACCAAAAAGTTTTTTAAAAATTTATGAA
GACAATAAAACTGTAGAGTCTGAGAAATTCCATTTGGATGTCGATATCTCTTATAAAGAA
ACAAAATAA
>vfdb~~~sell~~~NP_645577 (sell) staphylococcal enterotoxin L precursor
[Enterotoxin-like L (CVF065)] [Staphylococcus aureus subsp. aureus
MW2]
ATGAAAAAAAGATTATTATTTGTAATTGTTATTACTTTATTTATTTTTTCTTCTAATCAT
ACAGTCTTATCTAACGGCGATGTAGGTCCAGGAAACCTAAGAAATTTTTATACTAAATAT
GAATATGTGAATTTAAAGAATGTTAAAGACAAAAATTCACCAGAATCACACCGCTTAGAA
TACTCGTATAAAAATGATACATTGTATGCTGAATTTGACAATGAATATATAACTAGTGAT
CTAAAGGGAAAAAATGTCGATGTTTTTGGTATAAGCTATAAATATGGTTCTAACTCTCGT
ACTATATATGGTGGTGTTACTAAAGCAGAAAACAATAAATTAGATTCGCCAAGAATAATA
CCTATAAATTTAATTATCAATGGCAAGCATCAAACAGTTACAACTAAAAGTGTTTCTACA
GATAAAAAAATGGTTACCGCACAAGAAATAGATGTCAAACTAAGAAAATACTTGCAAGAT
GAATTTAATATTTATGGACACAATGATACTGGTAAAGGTAAAGAATACGGCACTTCTTCA
AAATTTTATAGCGGTTTTGATAAGGGGAGTGTAGTATTCCATATAAATGATGGTTCTAAT
TTCTCGTATGATTTATTTTACACAGGATACGGTCTTCCAGAAAGCTTCTTAAAAATTTAC
AAAGATAATAAAACTGTTGATTCAACACAATTTCATCTAGATGTCGAAATTTCAAAAAGA
TGA
>vfdb~~~selq~~~NP_646754 (selq) staphylococcal enterotoxin G precursor
[SE (VF0020)] [Staphylococcus aureus subsp. aureus MW2]
ATGAATAAAATATTTAGAGTTCTCACTGTTAGCTTGTTTTTCTTCACATTTTTAATAAAA
AACAATCTAGCATATGCTGATGTAGGGGTAATCAATCTTAGAAACTTTTATGCTAATTAT
CAACCTGAAAAGCTTCAAGGAGTTAGTTCTGGAAATTTTTCTACTTCTCATCAATTAGAG
TATATTGATGGAAAATACACTTTATATTCACAGTTTCATAATGAATATGAAGCGAAGAGA
TTAAAAGATCATAAAGTAGATATCTTTGGAATAAGTTACTCAGGTCTTTGTAATACAAAA
TATATGTATGGTGGAATTACGTTGGCAAATCAAAATTTAGATAAACCTAGAAATATACCT
ATTAATCTCTGGGTCAATGGTAAGCAAAATACTATATCTACAGACAAAGTTTCTACTCAA
AAAAAAGAAGTAACTGCTCAAGAGATTGATATTAAGTTACGAAAATACCTACAAAACGAA
TACAATATATATGGTTTTAATAAAACAAAGAAAGGTCAGGAATATGGATATAAGTCAAAA
TTTAATTCTGGGTTTAACAAAGGAAAAATTACTTTCCATTTAAATAATGAACCTTCTTTT
ACATACGATTTGTTTTACACCGGAACTGGTCAAGCAGAGAGTTTTTTAAAAATTTACAAT

GATAATAAAACTATAGATGCAGAGAATTTTCATTTGGATGTAGAGATTTCATATGAGAAA
ACTGAATAA
>vfdb~~~seg~~~LOY16_13225 (seg) phage associated @ Superantigen
enterotoxin SEG [SE (VF0020)] [Staphylococcus aureus AATYS]
atgaagaaattatctactgtaattattattttgattctagaaatagtttttcataatatg
aattatgtgaatgctcaacccgatcctaaattagacgaactaaataaagtaagtgattat
aaaaataataagggaactatgggtaatgtaatgaatctttatacgtctccacctgttgaa
ggaagaggagttattaattctagacagtttttatctcatgatttaatttttccaattgag
tataagagttataatgaggttaaaactgaattagaaaatacagaattagctaacaattat
aaagataaaaaagtagacattttggcgttccatattttatacatgtataatacctaaa
tctgaaccggatataaaccaaaattttggaggttgttgtatgtatggtggtcttacattt
aatagttcagaaaatgaaagagataaattaattactgtacaggtaacaatcgacaataga
caatcacttggatttacaataactacaaataagaatatggttactattcaggaactagat
tacaaagcaagacactggctcactaaagaaaaaaagctatacgagtttgatggttctgca
tttgaatctggatatataaaatttactgaaaagaacaatacaagtttttggtttgactta
tttcctaaaaaagaactagtaccttttgttccatataagtttttaaatatttacggagat
aataaagttgttgattctaagagtattaaaatggaagtatttcttaatactcactga
>vfdb~~~sen~~~UOO13789.1 (sen) staphylococcal enterotoxin type N [SE
(VF0020)] [Staphylococcus aureus ATCC 29213]
TTAATCTTTATATAAAAATACATCAATATGATAATTAGATGAGCTAACTGTTCTATTATCACTATAAAAT
TGAAAAAACTCTGCTCCCACTGAACCTTTTACGTTATATAAATCATAATAAAATGATTGATCTTGATGAT
TATGAGAATGAAAGAAAATGCATCCTTTTTGTATGTTACCGGTATCTTTATTGTATATTTTATATAAATT
TTCTAATTTAAATCGAACTTTAGTGTCTAATTCTTGTACTGTTACTTTAGCCTTTTTGGTTTTTATAACA
AAACCTTCTTGTTGGACACCATCTTTAAATACATTAACGCCTATAACTTTCTCTTCATCTAATTGATTTC
CATCATGTATCGTAACTCCTCCGTATAAGCATGATGTTTTTTCTTCAGTTAAGCCTACACATTTATTTCC
AAAATACAGTCCATAAATATCTATATTTTTTCCTTTAAATTGATTTGCTAAATCTGATGAGTTAAACTCA
ACTTTCAAACTAGAAGTTTTAAGTACGGATATATCAATATTTTTTAATATTATAGTATTATTCAGTAGTT
GATCTGTACTAATTTTATTTGACTCGTCTAATTGCCACGTTATATCAGTATAATAGCTTGTTAAATTAAA
TAACTTACTACTATCTAGATCAGATTTTTTCTTTAAATCTTTTTTGTCTACTTCAGCATTAACATAATTA
TTATTAATAAGACATAATAAAGTTATTATAATTGCAGCTATGTAGAACAATCTCAT
>vfdb~~~sei~~~UOO13790.1 (sei) staphylococcal enterotoxin type I [SE
(VF0020)] [Staphylococcus aureus ATCC 29213]
ATGAAAAAATTTAAATATAGTTTTATATTAGTTTTTATATTACTTTTTAACATTAAAGATCTTACGTATG
CTCAAGGTGATATTGGTGTAGGTAACTTAAGAAATTTCTATACAAAACATGATTATATAGATTTAAAAGG
CGTCACAGATAAAAACCTACCTATTGCAAATCAACTCGAATTTTCAACAGGTACCAATGATTTGATCTCA
GAATCTAATAATTGGGACGAAATAAGTAAATTTAAAGGAAAGAAACTGGATATTTTTGGCATTGATTATA
ATGGTCCTTGTAAATCTAAATACATGTATGGAGGGGCCACTTTATCAGGACAATACTTAAATTCTGCTAG
AAAAATCCCTATTAATCTTTGGGTTAATGGCAAACATAAAACAATTTCTACTGACAAAATAGCAACTAAT
AAAAAACTAGTAACAGCTCAAGAAATTGATGTTAAATTAAGAAGATATCTTCAAGAAGAATACAATATAT
ATGGTCATAATAACACTGGTAAAGGCAAAGAATATGGATATAAATCTAAATTTTATTCAGGTTTTAATAA
TGGGAAAGTTTTATTTCATTTAAATAATGAAAAATCATTTTCATATGATTTGTTTTATACAGGAGATGGA
CTGCCTGTAAGTTTTTTGAAAATTTATGAAGATAATAAAATAATAGAATCTGAAAAATTTCATCTTGATG
TCGAAATATCATATGTAGATAGTAACTAA
>vfdb~~~sem~~~UOO29694.1 (sem) staphylococcal enterotoxin type M [SE
(VF0020)] [Staphylococcus aureus TCH32767]
ATGAAAAGAATACTTATCATTGTTGTTTTATTGTTTTGCTATTCGCAAAATCATATCGCAACCGCTGATG
TCGGAGTTTTGAATCTTAGGAACTATTATGGTAGCTATCCAATTGAAGACCACCAAAGTATTAATCCTGA
AAATAATCATCTTTCGCATCAATTAGTTTTTTCTATGGATAATTCGACAGTAACAGCTGAATTTAAGAAC
GTTGATGATGTAAAGAAATTCAAAAATCATGCTGTAGATGTATATGGTCTAAGTTATAGTGGATATTGTT
TGAAAAACAAATATATATACGGTGGAGTTACATTAGCAGGTGATTATTTAGAGAAATCTAGACGTATTCC

TATTAATCTTTGGGTTAATGGAGAACATCAAACTATATCTACTGACAAAGTATCAACTAATAAAAAGTTA
GTAACAGCTCAAGAAATTGATACTAAATTAAGAAGATATCTACAAGAAGAATATAATATTTATGGCTTTA
ATGATACAAATAAAGGAAGAAATTATGGTAATAAGTCAAAATTTAGTTCTGGATTTAATGCAGGAAAAAT
ATTATTTCATTTGAATGATGGTTCATCATTTTCTTATGACTTATTTGATACTGGAACAGGACAAGCTGAA
AGTTTCTTAAAAATATATAATGACAACAAAACTGTCGAAACTGAAAAATTCCATTTAGATGTAGAAATAT
CTTATAAGGACGAAAGTTGA
>vfdb~~~seo~~~UOO31426.1 (seo) staphylococcal enterotoxin type O [SE
(VF0020)] [Staphylococcus aureus TCH32767]
ATTAAAAATAGTAAAGTAATGTTAAATGTATTATTATTAATTTTAAATTTAATTGCAATATGTAGTGTAA
ACAATGCATATGCAAATGAAGAAGATCCTAAAATAGAGAGTTTGTGTAAGAAGTCAAGTGTAGACCCTAT
TGCTTTACATAATATTAATGATGATTATATAAATAATCGATTTACGACAGTAAAATCAATTGTATCAACT
ACAGAAAAATTCTTAGACTTCGATTTATTATTTAAAAGTATTAATTGGTTAGATGGAATATCTGCTGAAT
TTAAAGATTTAAAAGTGGAATTTAGCTCATCAGCGATTTCTAAAGAATTTCTAGGAAAGACTGTTGATAT
TTATGGTGTTTACTATAAAGCACATTGTCATGGTGAGCATCAAGTGGATACTGCCTGTACATATGGTGGG
GTAACACCTCATGAAAATAATAAATTAAGCGAGCCTAAAAATATAGGAGTAGCTGTGTATAAGGATAATG
TAAATGTTAATACATTTATCGTTACTACAGATAAAAAGAAAGTTACTGCACAAGAACTTGATATTAAAGT
AAGAACAAAATTAAATAATGCATATAAATTGTATGACAGAATGACTAGTGATGTACAAAAAGGTTATATT
AAATTTCATTCTCATTCGGAGCATAAAGAATCATTTTATTATGATTTATTTTATATTAAAGGAAATTTAC
CAGATCAATATTTGCAAATTTATAATGATAATAAAACAATAGATTCATCAGACTATCATATTGATGTTTA
TTTATTTACATAA
>vfdb~~~sek~~~NMF17_04325 (sek) Superantigen enterotoxin SEK [SE
(VF0020)] [Staphylococcus aureus S136_1]
atgaaaaaattaatagttatcatattaataaatataataactttaagtgtctctaatagt
gccagcgctcaaggtgatataggaattgataatctcaggaatttttatacaaaaaaagac
tttgtagatttaaaagatgtaaaagacaatgatactcctatagctaatcaactacaattt
tcaaatgtatcttatgatttaatttcagaatcaaaagattttaataaatttagtaatttc
aagggaaaaaaacttgatgtttttggtattagttataatggccagtgtaacaataaatac
atatatggtggagtcacagctactaacgaatatctagataaacctagaaatatccctata
aatatatggatcaatggaaatcacaaaactatttctaccaataaagtttcgacaaataaa
aaatttgttaccgctcaagagattgatgtcaaattaagaaagtaccttcaagaagaatac
aacatttatggacataacggcactaaaaaaggagaagaatatggtcataaatcaaaattt
tattctgggtttaatattggtaaagtaacgttccatttaaataataatgacactttttca
tatgatttattctacacaggagatgatgggttaccaaaaagtttttttaaaaatttacgaa
gacaataaaactgtagagtctgagaaattccatttggatgtcgatatttcttataaagaa
acgatataa
>vfdb~~~seq~~~NMF17_04330 (seq) staphylococcal enterotoxin type Q [SE
(VF0020)] [Staphylococcus aureus S136_1]
atgaataaaatatttagagtactcactgttagcttgtttttcttcacatttttaataaaa
aacaatctagcatatgctgatgtaggggtaatcaaccttagaaacttttatactaattat
caaccagaaacgcttcaaggagttagttctggaaatttttctacttctcatcaattagag
tatattgatggaaaatacactttatattcacagtttcataatgaatatgaagcgaagaga
ttaaaagatcataaagtagatatctttggaataagttactcaggtctttgtaatacaaaa
tatatgtatggcggaattacgttggcgaatcaaaatttagataaacctagaaatatacct
attaatctctgggtcaatggtaagcaaaacactatatctacagacgaggtttctactcaa
aaaaagaggtaactgctcaagagattgatattaagttacgaaaatacctacaaaacgaat
acaatatatatggttttaataaaacaaaaaaaggtcaggaatatggatatcagtcaaaat
ttaattctggatttaacaaaggaaaaattactttccatttaaataatgaaacttcttta
catacgatttgttttacaccggaactggtcaagcagagagtttttttaaaaatttacaatg
ataataaaactatagatgcagagaattttcatttggatgtagagatttcatatgagaaga
ctgaataa

>vfdb~~~sel~~~SAMEA2236454_00292 (sel) Exotoxin, phage associated @ Superantigen enterotoxin SEL [SE (VF0020)] [Staphylococcus aureus S136_1]
atgaaaaaaagattattatttgtaattgttattactttatttatttttttcttctaatcat
acagtcttatctaacggcgatgtaggtacaggaaacctaagaaatttttatactaaatat
gaatatgtgaatttaaagaatgttaaagacaaaaattcaccagaatcacaccgcttagaa
tactcgtataaaaatgatacattgtatgctgaatttgacaatgaatatataactagtgat
ctaaagggaaaaaatgtcgatgtttttggtataagctataaatatggttctaactctcgt
actatatatggtggtgttactaaagcagaaaacaataaattagattcgccaagaataata
cctataaatttaattatcaatggcaagcatcaaacagttacaactaaaagtgtttctaca
gataaaaaaatggttaccgcacaagaaatagatgtcaaactaagaaaatacttgcaagat
gaatttaatatttatggacacaatgatactggtaaaggtaaagaatacggcacttcttca
aaatttatagcggttttgataaggggagtgtagtatttcatatgaatgatggttctaat
ttctcgtatgatttattttacacaggatacggtcttccagaaagcttcttaaaaatttac
aaagataataaaactgttggttcaacacaatttcatctagatgtcgaaatttcaaaaaga
tga
>vfdb~~~se1y~~~BAS21358.1 (sey) enterotoxin Y [SE (VF0020)] [Staphylococcus aureus AV3008]
ATGAAAGCGAAACTATGGTTTTTATTGACAACTTTGGCATTTTTGATTGCAGTAACGGAATCTATTGGAA
TAGCAGAAGTAAAAGCAAAAACAACTGGATTGATTACAGAAAATAGTAATGACAGTTTAAAAGAGCATTA
TGCACAAAAATTTGAAGTTTATACGAATAAAGAAGTAACAGGAGTTGGGGAGAATTATATAGACACGAAA
GTTGACACCTACAATGTACGGACAGTGCTCTACAACACTGATTATTTGAAACAGTTTAAAAATCAAGACA
AAGTTAATATATGGGGAACATTATATGAAAACCAACAATCTAAAGTATATCGTGGCACAGTAGTTAAGTA
TGATCCTATATCAAAAGTAACTAACTTATCTTATAGAATGAACTTGTTTGTTAACGGTCATCAAACTAAA
GTGAATCCAGACAGTTTATTAGAAGTTAAGAATAAACAGATTTCTCTAAAAGAAACAGATTTTAGAATTA
GAAAATATTTATTAGAAAAAGAACACCTATATAGTAATTACAATTCAGGGGAACTAATTATAGAAATGAA
AAATGGAGCAAGACATAAAATAGATTTAGGCGACATATTGAGTGACTCACAAGAAAAAACGTTTGATTTT
GATAATATTAGTCATATAGATATTTATATGAAATAG
>vfdb~~~rpoB~~~SAUSA300_0527 (rpoB) RNA polymerase beta subunit [SE (VF0020)] [Staphylococcus aureus FPR3757]
ttggcaggtcaagttgtccaatatggaagacatcgtaaacgtagaaactacgcgagaatt
tcagaagtattagaattaccaaacttaatagaaattcaaactaaatcttacgagtggttc
ctaagagaaggtttaatcgaaatgtttagagacatttctccaattgaagattttactggt
aatttgtcattagagtttgtggattaccgtttaggagaaccaaaatatgatttagaagaa
tctaaaaaccgtgacgctacttatgctgcacctcttcgtgtaaaagtgcgtctaatcatt
aaagaaacaggagaagttaaagaacaagaagtctttatgggtgatttcccattaatgact
gatacaggtacgttcgttatcaatggtgcagaacgtgtaatcgtatctcaattagttcgt
tcaccatccgtttatttcaatgaaaaaatcgacaaaaaatggtcgtgaaaactatgatgca
acaattattccaaaccgtggtgcatggttagaatatgaaacagatgctaaagatgttgta
tacgtacgtattgatagaacacgtaaactaccattaacagtattgttacgtgcattaggt
ttctcaagcgaccaagaaattgttgaccttttaggtgacaatgaatatttacgtaatact
ttagagaaagacggcactgaaaacactgaacaagcgttattagaaatctatgaacgttta
cgtccaggtgaaccaccaactgttgaaaatgctaaaagtctattgtattcacgtttcttt
gatccaaaacgctatgacttagcaagcgtgggtcgttataaaacaaacaaaaaattacat
ttaaaacatcgtttatttaatcaaaaattagctgagccaattgtaaatactgaaactggt
gaaattgtagttgaagaaggtacagtgcttgatcgtcgtaaaatcgacgaaatcatggat
gtacttgaatcaaatgcaaacagcgaagtgtttgaattgcatggtagcgttatagacgag
ccagtagaaattcaatcaattaaagtatatgttcctaacgatgatgaaggtcgtacgaca
actgtaattggtaatgctttccctgactcagaagttaaatgcattacaccagcagatatc
attgcttcaatgagttacttctttaacttattaagcggtattggatatacagatgatatt

gaccatttaggtaaccgtcgtttacgttctgtaggtgaattactacaaaaccaattccgt
atcggtttatcaagaatggaaagagttgtacgtgaaagaatgtcaattcaagatactgag
tctatcacacctcaacaattaattaatattcgacctgttattgcatctattaaagaattc
tttggtagctctcaattatcacaattcatggaccaagcaaacccattagctgagttaacg
cataaacgtcgtctatcagcattaggacctggtggtttaacacgtgaacgtgctcaaatg
gaagtacgtgacgttcactactctcactatggccgtatgtgtccaattgaaacacctgag
ggaccaaacattggattgattaactcattatcaagttatgcacgtgtaaatgaattcggc
tttattgaaacaccatatcgtaaagttgatttagatacacatgctatcactgatcaaatt
gactatttaacagctgacgaagaagatagctatgttgtagcacaagcaaactctaaatta
gatgaaaatggtcgtttcatggatgatgaagttgtatgtcgtttccgtggtaacaataca
gttatggctaaagaaaaaatggattatatggatgtatcgccgaagcaagttgtttcagca
gcgacagcatgtattccattcttagaaaatgatgactcaaaccgtgcattgatgggtgcg
aacatgcaacgtcaagcagtgcctttgatgaatccagaagcaccatttgttggtacaggt
atggaacacgttgcagcacgtgattctggtgcggctattacagctaagcacagaggtcgt
gttgaacatgttgaatctaatgaaattcttgttcgtcgtctagttgaagagaacggcgtt
gagcatgaaggtgaattagatcgctatccattagctaaatttaaacgttcaaactcaggt
acatgttacaaccaacgtccaatcgttgcagttggagatgttgttgagtataacgagatt
ttagcagatggaccatctatggaattaggagaaatggcattaggtagaaacgtagtagtt
ggtttcatgacttgggacggttacaactatgaggatgccgttatcatgagtgaaagactt
gtgaaagatgacgtgtatacttctattcatattgaagagtatgaatcagaagcacgtgat
actaagttaggacctgaagaaatcacaagagatattcctaatgtttctgaaagtgcactt
aagaacttagacgatcgtggtatcgtttatattggtgcagaagtaaaagatggagatatt
ttagttggtaaagtaacgcctaaaggtgtaactgagttaactgccgaagaaagattgtta
catgcaatctttggtgaaaaagcacgtgaagttagagatacttcattacgtgtacctcac
ggcgctggcggtatcgttcttgatgtaaaagtattcaatcgtgaagaaggcgacgataca
ttatcacctggtgtaaaccaattagtacgtgtatatatcgttcaaaaacgtaaaattcat
gttggtgataagatgtgtggtcgacatggtaacaaaggtgtcatttctaagattgttcct
gaagaagatatgccttacttaccagatggacgtccgatcgatatcatgttaaatcctctt
ggtgtaccatctcgtatgaacatcggacaagtattagagctacacttaggtatggctgct
aaaaatcttggtattcacgttgcatcaccagtatttgacggtgcaaacgatgacgatgta
tggtcaacaattgaagaagctggtatggctcgtgatggtaaaactgtactttatgatgga
cgtacaggtgaaccattcgataaccgtatttcagtaggtgtaatgtacatgttgaaactt
gcgcacatggttgatgataaattacatgcgcgttcaacaggaccatattcacttgttaca
caacaaccacttggcggtaaagcgcaattcggtggacaacgttttggtgagatggaggta
tgggcacttgaagcatatggtgctgcatacacattacaagaaatcttaacttacaaatcc
gatgatacagtaggacgtgtgaaaacatacgaggctattgttaaaggtgaaaacatctct
agaccaagtgttccagaatcattccgagtattgatgaaagaattacaaagtttaggttta
gatgtaaaagttatggatgagcaagataatgaaatcgaaatgacagacgttgatgacgat
gatgttgtagaacgcaaagtagatttacaacaaaatgatgctcctgaaacacaaaaagaa
gttactgattaa

Supp Table 1

| | Marker | Color/Format |
|---|---|---|
| Flow | CD24 | APC |
| Flow | CD45 | APC-eFluor 780 |
| Flow | CD11b | Brilliant Blue 515 |
| Flow | Ly-6G | Brilliant Ultraviolet 395 |
| Flow | CD11c | Brilliant Ultraviolet 615 |
| Flow | CD103 | Brilliant Ultraviolet 737 |
| Flow | Ly-6C | Brilliant Violet 570 |
| Flow | CD169 | Brilliant Violet 605 |
| Flow | Siglec-F | Brilliant Violet 750 |
| Flow | F4/80 | eFluor 450 |
| Flow | HLA DR | PE |
| Flow | CD125 | PE-Cy7 |
| Flow | CX3CR1 | PE-Fire 810 |
| Flow | CD193 | PerCP-Cy5.5 |
| Flow | | |
| Flow | CD64 | Brilliant Violet 650 |
| Flow | Live/Dead Fix Blue | Live/Dead Fix Blue |
| Flow | CD8 | BUV737 |
| Flow | CD4 | BV650 |
| Flow | CD3 | NovaFluor Blue 660-120S |
| Flow | TCRgd | PE |
| Flow | NKg2D | PE-Cy7 |
| Flow | CD45 | APC-eFluor 780 |
| Flow | NK1.1 | Alexa Fluor 700 |
| Flow | TCR Vb8.1 8.2 | BUV615 |
| Flow | TCR Vb6 | BV510 |
| Flow | NKp46 | PerCP-eFluor 710 |
| Flow | Live/Dead Far Red | |
| IHC | CD3 | |
| IHC | CD4 | |
| IHC | CD8 | |
| IHC | CD19 | |

| | |
|---|---|
| IHC | Anti-MBP |
| In vivo | anti-IFN-g antibodies |

| Clone |
| --- |
| 30-F1 |
| 30-F11 |
| M1/70 |
| 1A8-Ly6g |
| N418 |
| M290 |
| HK1.4 |
| 3D6.112 |
| E50-2440 |
| BM8 |
| DIH37 |
| SA011F11 |
| J073E5 |
| 53-6.7 |
| RM4-5 |
| 145-2C11 |
| GL3 |
| CX5 |
| PK136 |
| KJ16-33 |
| RR4-7 |
| 29A1.4 |
| Clone E4T1B |
| Clone 4SM95 |
| Clone 4SM1 |
| Clone D4V4B |

| |
|---|
| clone MT2-14.7.3 |
| Clone H22 |

| Company |
| --- |
| BD Biosciences [877.232.8995] |
| Thermo Fisher Scientific [1 800 955 6288] |
| BD Biosciences [877.232.8995] |
| Thermo Fisher Scientific [1 800 955 6288] |
| Thermo Fisher Scientific [1 800 955 6288] |
| BD Biosciences [877.232.8995] |
| BioLegend [877-246-5343] |
| BioLegend [877-246-5343] |
| BD Biosciences [877.232.8995] |
| Thermo Fisher Scientific [1 800 955 6288] |
| BioLegend [877-246-5343] |
| BioLegend [877-246-5343] |
| BioLegend [877-246-5343] |
| BioLegend [877-246-5343] |
| BD Biosciences [877.232.8995] |
| Thermo Fisher Scientific [1 800 955 6288] |
| Thermo Fisher |
| Thermo Fisher |
| Thermo Fisher |
| Thermo Fisher |
| Thermo Fisher |
| Thermo Fisher |
| eBio |
| BD |
| BD |
| eBio |
| Thermo Fisher |
| Cell Signaling |
| Invitrogen |
| Invitrogen |
| Cell Signalling |

| |
| --- |
| Mayo Clinic, AZ |
| BioXcell |
